# Barcoded Rabies In Situ Connectomics for high-throughput reconstruction of neural circuits

**DOI:** 10.1101/2025.07.16.665048

**Authors:** Alexander Becalick, Antonin Blot, Molly Strom, Petr Znamenskiy

## Abstract

Sequencing of oligonucleotide barcodes holds promise as a high-throughput approach for reconstructing synaptic connectivity at scale (1). Rabies viruses can act as a vehicle for barcode transmission, thanks to their ability to spread between synaptically connected cells (2, 3). However, applying barcoded rabies viruses to map synaptic connections *in vivo* has proved challenging (4–7). Here, we develop Barcoded Rabies In Situ Connectomics (BRISC) for high-throughput connectivity mapping in the mouse brain. To ensure that the majority of post-synaptic “starter” neurons are uniquely labeled with distinct barcode sequences, we first generated libraries of rabies viruses with sufficient diversity to label >1000 neurons uniquely. To minimize the probability of barcode transmission between starter neurons, we developed a strategy to tightly control their density. We then applied BRISC to map inputs of single neurons in the primary visual cortex (V1). Using *in situ* sequencing, we read out the expression of viral barcodes in rabies-infected neurons, while preserving spatial information. We then matched barcode sequences between starter and presynaptic neurons, mapping the inputs of 385 neurons and identifying 7,814 putative synaptic connections. The resulting connectivity matrix revealed layer- and cell-type-specific local connectivity rules and topographic organization of long-range inputs to V1. These results show that BRISC can simultaneously resolve the synaptic connectivity of hundreds of neurons while preserving spatial information, enabling reconstruction of neural circuits at an unprecedented scale.

## Introduction

Mechanistic understanding of how networks of neurons carry out neural computations requires knowledge of their underlying synaptic connectivity. However, current methods for reconstructing synaptic connections are extremely laborious and typically limited to mapping connections between nearby neurons. Although recent years have seen dramatic advances in connectomics approaches based on volume electron microscopy, they remain limited to reconstructing single volumes up to ∼ 1 mm^3^ (8–10). The effort required to process a single volume makes it impractical to compare connectivity rules between different areas or experimental conditions, such as in models of psychiatric or neurological diseases. Adding to the challenge, recent transcriptomic studies have revealed a surprising diversity of cell types throughout the brain (11–15), raising speculation that different transcriptomic cell types may subserve distinct computational roles owing to their differential connectivity (16, 17). Therefore, there is an acute need for methods that can read out synaptic connectivity with high throughput while preserving molecular information.

Over a decade ago, Zador and colleagues (1) proposed that synaptic connectivity could be read out at a massive scale using DNA sequencing. By labeling individual neurons with unique oligonucleotide “barcodes” and having them transmit these barcodes to their synaptic partners, the connectivity matrix could be read out by matching barcode sequences across neurons. Owing to its scalability, this approach has the potential to revolutionize connectomics, but it has been challenging to implement for mapping connectivity *in vivo* (4, 5, 18, 19).

Rabies viruses can propagate transynaptically from infected neurons to their presynaptic partners and have been proposed as a potential vehicle for barcode transmission (4–7). Conventional monosynaptic rabies tracing labels inputs to populations of interest using deletion mutant rabies viruses that lack the gene encoding rabies glycoprotein (G), which is essential for the generation of infectious virions (2, 3). Therefore, only cell populations where G expression is provided *in trans*, referred to as *starter cells*, can transmit the virus to their direct presynaptic partners (3). As long as presynaptic cells do not express G themselves, the virus cannot propagate further, leading to monosynaptically restricted tracing.

Barcoded rabies tracing aims to map inputs to many starter neurons simultaneously while maintaining single-cell resolution not afforded by traditional bulk rabies tracing. By using rabies viruses that express random mRNA barcodes, connections between individual presynaptic and starter neurons are identified by matching their barcode sequences (Fig. 1a). Despite the promise of this technology, barcoded rabies tracing has not yet been applied to map synaptic connectivity at scale. An essential prerequisite for applying such an approach is the unique labeling of starter neurons with distinct barcode sequences. However, this has proved difficult to achieve in practice (4, 5). First, multiple starter cells may be infected by rabies virions carrying the same sequence during the initial viral injection. Second, non-unique labeling can also occur if starter cells transmit barcodes to one another. Both of these factors have hampered previous efforts to map connectivity using barcoded rabies viruses (5). Although the generation of higher diversity libraries has since been reported, they have not yet been applied *in vivo* (7).

**Fig. 1.**
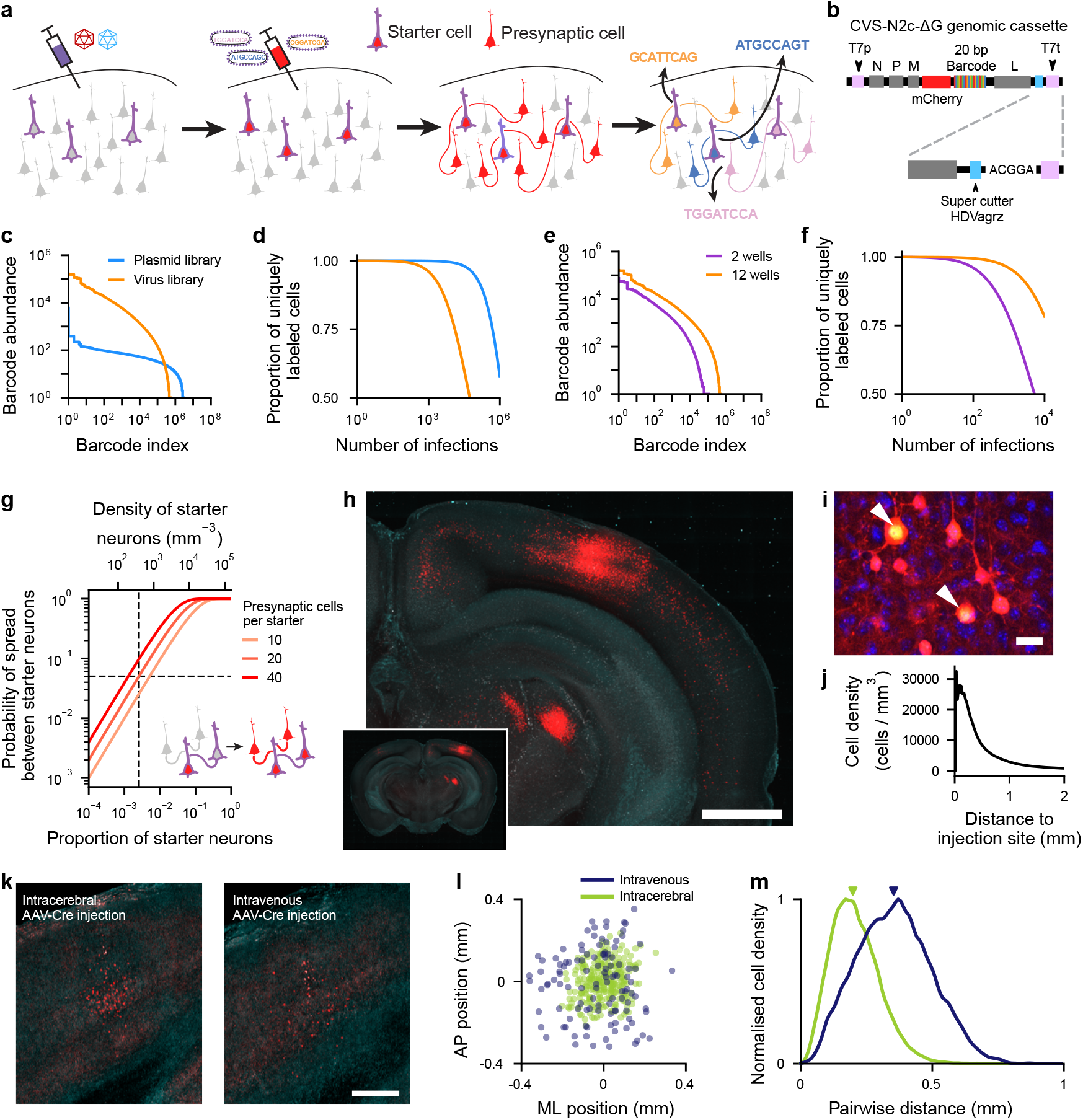
Generation and delivery of barcoded rabies virus libraries. **a**, BRISC experimental outline. **b**, Barcoded rabies virus genome plasmid schematic. The barcode was integrated in the 3’ UTR of the mCherry transgene inserted in place of the G glycoprotein gene. The HDVagrz ribozyme was replaced with a supercutter version to ensure the correct 3’ terminus of the rabies genome transcript. **c**, Barcode abundance distribution of plasmid and viral libraries. Barcodes are sorted according to their abundance in the sequencing library, estimated using unique molecular identifiers (see Methods). **d**, Proportion of uniquely labeled starter neurons as a function of the number of independent infections. The barcoded viral library has sufficient complexity to label >1,000 starter cells. **e-f**, Library complexity and the number of neurons that can be uniquely labeled with a library depend on the library production scale. **g**, Predicted relationship between the density of starter neurons and the frequency of barcode transmission between them. For 20 presynaptic cells per starter, the 5% probability of spread between starters (horizontal dashed line) corresponds to 384 cells per mm^3^ (vertical dashed line). **h**, mCherry fluorescence in a section of mouse brain infected with the barcoded rabies virus library in the primary visual cortex, imaged using serial two-photon tomography. Maximum intensity projection over 0.5 mm. Scale bar – 1 mm. **i**, Identification of starter neurons based on nuclear mCherry fluorescence in a confocal microscopy image (maximum intensity projection spanning 4.8 µm, blue – DAPI, red and green – mCherry channel with different contrast limits, see Methods). Scale bar – 20 µm. **j**, Density of rabies-infected neurons as a function of distance to the injection site. **k**, mCherry fluorescence in a section of mouse visual cortex infected locally with Cre-dependent AAV helper virus and either an intracerebral or intravenous injection of AAV-Cre. **l**, Spatial distribution of detected mCherry cells from intracerebral or intravenous AAV-Cre injections. **m**, Kernel density estimates (KDEs) of the distribution of pairwise distances between mCherry-positive cells in intracerebral and intravenous AAV-Cre injections, median distances (triangles) – 0.20 mm, 256 cells, and 0.35 mm, 114 cells respectively.

Here, we describe the development of Barcoded Rabies In Situ Connectomics (BRISC), which overcomes these obstacles to map putative presynaptic ensembles of hundreds of starter neurons simultaneously. BRISC uses libraries of rabies viruses with sufficient diversity to uniquely label >1,000 starter neurons while ensuring that >95% of them carry unique barcodes. To minimize the frequency of transmission of barcodes between starter neurons, their density is tightly controlled by limiting dilution of an AAV expressing Cre-recombinase. In contrast to Cre delivery targeting specific transcriptionally or anatomically defined cell types (5), the stochastic infection of starter neurons by AAV-Cre enables BRISC to map inputs of diverse cell populations simultaneously. After allowing the rabies virus to spread from starter to presynaptic neurons, barcodes are read out from intact tissue sections using *in situ* sequencing, preserving spatial information.

We applied BRISC to map monosynaptic inputs onto single neurons in the mouse primary visual cortex (V1). First, by reconstructing local connectivity, we identified layer-specific connectivity rules that are consistent with the established wiring patterns of V1. Second, by combining BRISC with *in situ* sequencing of cell-type marker transcripts, we demonstrate its potential to relate the connectivity of individual neurons to their molecular identity. Finally, by applying BRISC to map the spatial distribution of long-range inputs, we reveal the topographic organization of corticocortical inputs onto single V1 neurons. Together, these results demonstrate the potential of BRISC for high-throughput reconstruction of both long-range and local microcircuits.

## Results

### Generating high diversity barcoded rabies libraries

Identification of individual synaptic connections using barcoded rabies tracing depends on the unique labeling of starter neurons by distinct barcode sequences. We chose the CVS-N2c(ΔG) strain of rabies virus as the basis for generating diverse barcoded viral libraries. Compared to the widely used SAD-B19(ΔG) strain, CVS-N2c(ΔG) shows enhanced transsynaptic spread and lower neuronal cytotoxicity (20). We first optimized the production of high-diversity barcoded rabies libraries. During the packaging of glycoprotein-deficient rabies viruses, virions are first generated by rescue from plasmid-encoded viral cDNA. This rescue is highly inefficient (21) and therefore constitutes the primary bottleneck reducing library diversity (4). The first step of rabies viral rescue is the transcription of the viral antigenome from the transfected plasmid DNA. The efficiency of rescue is increased by producing the correct 5’ and 3’ terminal sequences of the transcribed antigenome, which are generated by ribozyme-mediated cleavage of the transcript ends (22). Previously described plasmids for rescue of CVS-N2c(ΔG) contain an incomplete sequence of the hepatitis delta virus antigenomic ribozyme (HDVagrz) responsible for generating the 3’ end of the antigenome (23). To improve rescue efficiency, we first inserted the full-length HDVagrz in the CVS-N2c(ΔG) genome plasmid. Then we inserted a 20-nucleotide random barcode sequence downstream of an mCherry transgene in the CVS-N2c(ΔG) genome by inverse PCR (Fig. 1b).

We used the resulting plasmid library to package barcoded rabies viruses and optimized the viral packaging protocol to improve the diversity of the viral library (see Methods). We quantified the diversity of the plasmid and the resulting viral libraries using Illumina sequencing (Fig. 1c; Supplementary Fig. 1a,b). As a consequence of the bottleneck imposed by the inefficiency of rabies virus rescue, the resulting viral library contained five-fold fewer unique barcodes than the plasmid library (2.8 × 10^6^ barcodes in the plasmid library compared to 5.3 × 10^5^ in the viral library). Furthermore, while the 100 most abundant barcodes accounted for only 0.06% of the plasmid library, they comprised 3.9% of the viral library. These results demonstrate that the diversity of barcoded virus libraries is constrained by both the inefficiency of viral rescue and the uneven amplification of viral barcodes during subsequent viral production steps.

We next calculated how many neurons could be labeled uniquely if barcodes were randomly sampled from each library based on their abundance distributions (Fig. 1d) (24). The plasmid library had sufficient diversity to label >80,000 cells while ensuring that 95% of them received unique barcodes. The same threshold was reached at 1,320 cells for the viral library. Therefore, the resulting barcoded viral libraries have sufficient diversity to trace inputs of >1,000 starter neurons despite the reduction in barcode diversity during viral packaging, surpassing most previously described barcoded rabies libraries (Supplementary Fig. 1a,b) (4, 5, 7, 25).

We next asked how the diversity of the rabies viral library depends on the scale of the viral rescue. Compared to a library generated from transfection of 12 wells, a library generated from 2 wells had a substantially lower diversity, sufficient to label only ∼ 132 neurons with the same 95% unique threshold (Fig. 1e-f). This indicates that viral rescue constitutes a key bottleneck in the production of barcoded rabies viruses, suggesting that library diversity could be further improved by increasing its scale.

### Controlling starter neuron density

Non-unique labeling of starter neurons can also result from the transsynaptic spread of viral barcodes between starters. Under the simplifying assumption of random connectivity, the probability that a given starter neuron will transmit its barcode to another starter cell is determined by the number of neurons transsynaptically infected by each starter cell, and by the proportion of neurons at the injection site that are starter cells (Fig. 1g, see Methods). To determine the average number of nearby cells labeled by each starter neuron, we injected starter AAVs FLEX-H2B-mCherry-P2A-N2cG and FLEX-H2B-EBFP2-P2A-TVA in the mouse primary visual cortex (V1) alongside low-titer AAV hSyn-Cre to control starter cell density, followed by CVS-N2c(ΔG) mCherry barcoded rabies virus (Fig. 1h). We then quantified the distribution of rabies-infected cells throughout the brain and the number of starter neurons at the injection site. Starter neurons were identified based on nuclear fluorescence of the H2B-mCherry-P2A-N2cG construct (Fig. 1i, see Methods). On average, we detected 66.6 presynaptic neurons per starter (17,505 presynaptic neurons for 263 starter cells). As the expression of starter AAVs is restricted to the injection site, local but not long-range transsynaptic spread can lead to the transmission of barcodes between starter neurons. Therefore, we quantified the density of rabies-infected neurons as a function of distance from the injection site (Fig. 1j). As expected, the density was highest near the center of the injection site, with each starter neuron infecting an average of 20.9 presynaptic cells within 0.5 mm of the injection center. Together, these observations suggest that to limit the proportion of starter neurons that transmit their barcodes to other starter cells to <5%, the proportion of starter cells at the injection site must be limited to 2.6 × 10^-3^, which corresponds to a density of ∼ 400 starter cells per mm^3^ (Fig. 1g, see Methods).

As the probability of transsynaptic spread rapidly drops off with distance, in an ideal scenario starter neurons would be uniformly distributed throughout the brain region of interest. However, stereotactic injection of starter AAVs inevitably results in the clustering of starter neurons. This would lead to a high local density of starter cells, thereby increasing the probability of barcode transmission between them. To control the density of starter neurons while minimizing their spatial clustering, we used tail vein injections of blood-brain-barrier penetrating AAV PHP.eB hSyn-Cre (26). Intravenous delivery resulted in a more uniform distribution of starter cells compared to local intracerebral injection (Fig. 1k-m). Consequently, the pairwise distances between starter cells were higher after intravenous delivery of Cre compared to intracerebral delivery (Fig. 1m).

Finally, to determine the conditions necessary to achieve the desired density of starter neurons, we injected serial dilutions of AAV PHP.eB hSyn-Cre into tdTomato Cre reporter mice and quantified the number of tdTomato-positive neurons throughout the brain using serial two-photon tomography. The density of cells expressing Cre scaled with increasing titers of AAV (Supplementary Fig. 1c). Therefore, systemic delivery of AAV PHP.eB hSyn-Cre provides a means to control the density of starter neurons in order to limit the probability of transsynaptic spread between them.

### *In situ* sequencing of rabies viral barcodes

We next applied BRISC to systematically map synaptic inputs of V1 neurons. We used a tail vein injection of AAV PHP.eB hSyn-Cre and stereotactic injections of AAVs FLEX-H2B-mCherry-P2A-N2cG and FLEX-H2B-EBFP2-P2A-TVA to induce sparse expression of TVA and N2cG from starter AAVs. We made three injections arranged along the mediolateral extent of the primary visual cortex, allowing us to sample starter neurons across different layers, cell types, and retinotopic coordinates of V1. Thirteen days later, we injected the barcoded rabies virus library described in Fig. 1c at the same three injection sites as the AAVs. We allowed the barcodes to spread to presynaptic neurons for seven days before collecting the brain tissue (Fig. 2a).

**Fig. 2.**
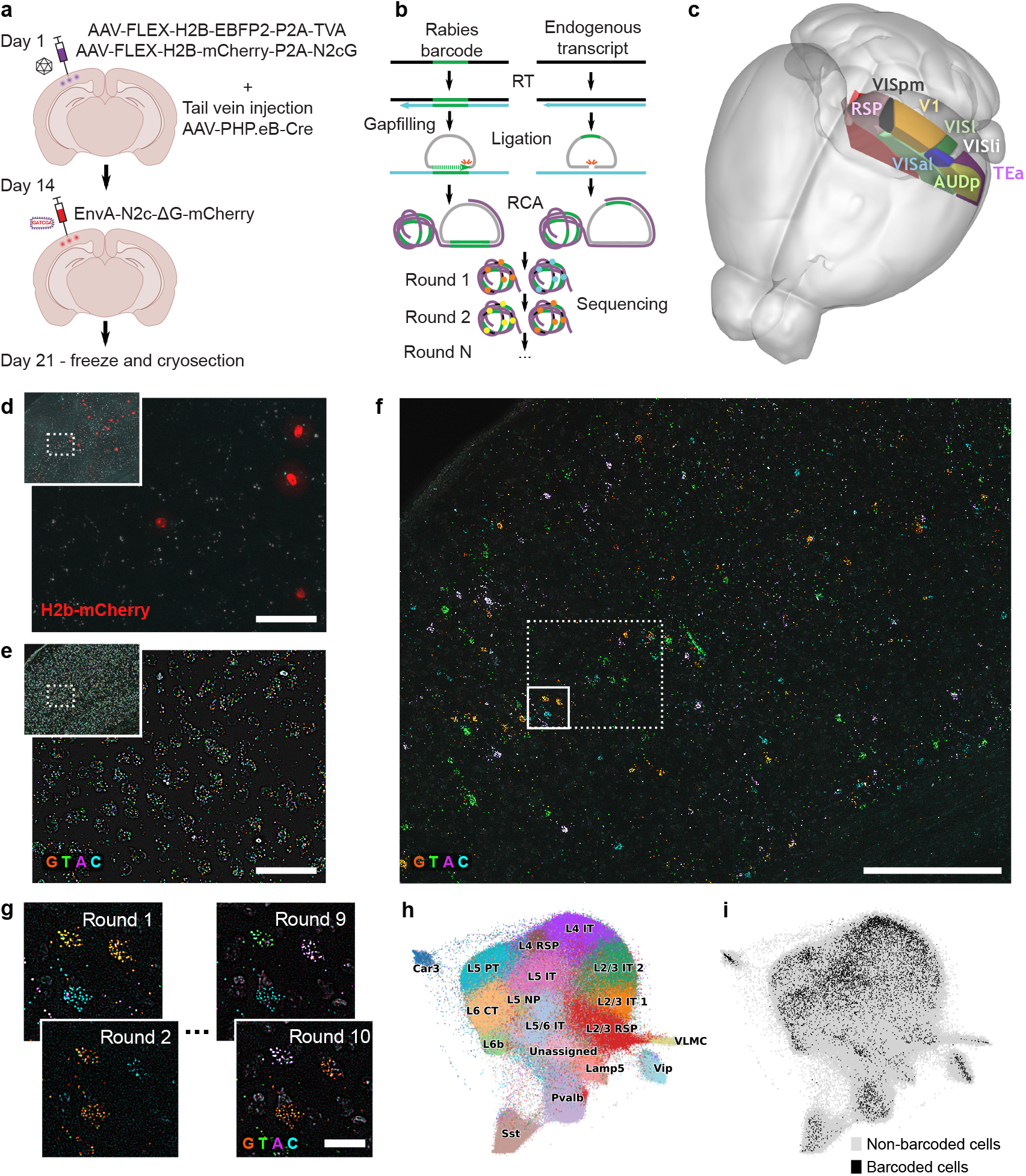
Sequencing rabies virus barcodes *in situ*. **a**, Virus injection timeline. Three injections were arranged along the mediolateral extent of primary visual cortex at the same anterior-posterior location. **b**, Schematic of the protocol for sequencing rabies virus barcodes alongside cell type marker transcripts. **c**, Extent of the *in situ* sequencing volume in the mouse brain. **d**, Nuclear mCherry fluorescence identifies starter neurons expressing G glycoprotein. Scale bar – 50 µm. **e**, Example sequencing round during *in situ* sequencing of cell type marker transcripts. Scale bar – 50 µm. **f**, Example sequencing round of rabies virus barcodes. Dashed and solid rectangles indicate the image regions depicted in **d**,**e** and **g**, respectively. Scale bar – 250 µm. **g**, Three example neurons infected with barcoded rabies virus across sequencing rounds. Scale bar – 20 µm. **h**, UMAP representation of cell type marker gene expression across cells, color-coded according to their assigned cell type. **i**, Distribution of rabies-infected cells in UMAP space.

Ensuring that starter neurons are uniquely labeled requires dense sequencing to capture the majority of starter neurons, precluding the use of single-cell or single-nucleus RNA sequencing, which recovers only a subset of infected neurons (4, 5). To overcome this, we developed a strategy for sequencing rabies virus barcodes *in situ* alongside endogenous transcripts for cell type identification based on BARseq2 and coppaFISH (see Methods) (27, 28). This method uses rolling circle amplification followed by sequencing-by-synthesis from intact tissue sections (Fig. 2b), which is performed by imaging the tissue over multiple sequencing rounds to read out barcode sequences. Recovering all starter neurons in a barcoded rabies tracing experiment by *in situ* sequencing requires imaging tens of tissue sections, which limits the throughput of barcoded rabies tracing experiments. To increase the speed of data acquisition, we designed custom sequencing flowcells and an imaging setup based on a conventional epifluorescence microscope, which allowed for the parallel imaging of the four fluorophores associated with each DNA base. This enabled the simultaneous sequencing of up to 40 coronal sections of mouse brain (Supplementary Fig. 2). We processed serial cryosections spanning 800 µm of brain tissue along the anterior-posterior axis, covering a total imaging volume of ∼ 12 mm^3^ (Fig. 2c). We sequenced 10 nucleotides of the viral barcode as the number of starter cells that could be uniquely identified plateaued at that length (Supplementary Fig. 3, see Methods).

Nuclear localized H2B-mCherry expressed from AAV FLEX-H2B-mCherry-P2A-N2cG is preserved in cryosections, allowing the detection of starter neurons without any further staining (Fig. 2d). The sequenced volume contained 92% of all mCherry-positive cells across the injection site (Supplementary Fig. 4a, see Methods). Based on the distribution of presynaptic cells in bulk tracing experiments (Fig. 1h-j), we estimated that the sequenced volume included approximately 44% of all presynaptic cells across the brain (Supplementary Fig. 4b). We then sequenced the transcripts of a panel of cell-type marker genes (Fig. 2e, Supplementary Fig. 4c), selected to facilitate the classification of V1 cell types, followed by the rabies virus barcode (Fig. 2f-g). Finally, we used *in situ* hybridization to measure the expression of highly expressed cell-type marker genes, followed by DAPI staining to assist with cell segmentation. Barcodes were assigned to cells using a probabilistic algorithm that relies on spatial clustering of barcodes near the infected cell nucleus (see Methods). We used the expression of cell-type marker transcripts to classify cells into major cortical cell types (Fig. 2h). Rabies-infected cells were intermingled with uninfected cells in UMAP space and were assigned to clusters corresponding to major cell types of V1 (Fig. 2i), indicating that rabies infection did not interfere with cell type identification.

We classified rabies-infected neurons as starter cells if they expressed H2B-mCherry or as presynaptic cells if they did not, and examined the barcode sequences present in individual cells. The majority (93%) of rabies-infected neurons contained only one barcode (Fig. 3a), although starter cells were more likely to contain multiple barcodes than presynaptic cells. This finding is consistent with previous studies (4, 6) and is likely a consequence of multiple EnvA/TVA-mediated infection events occurring per cell in starter neurons after viral injection.

**Fig. 3.**
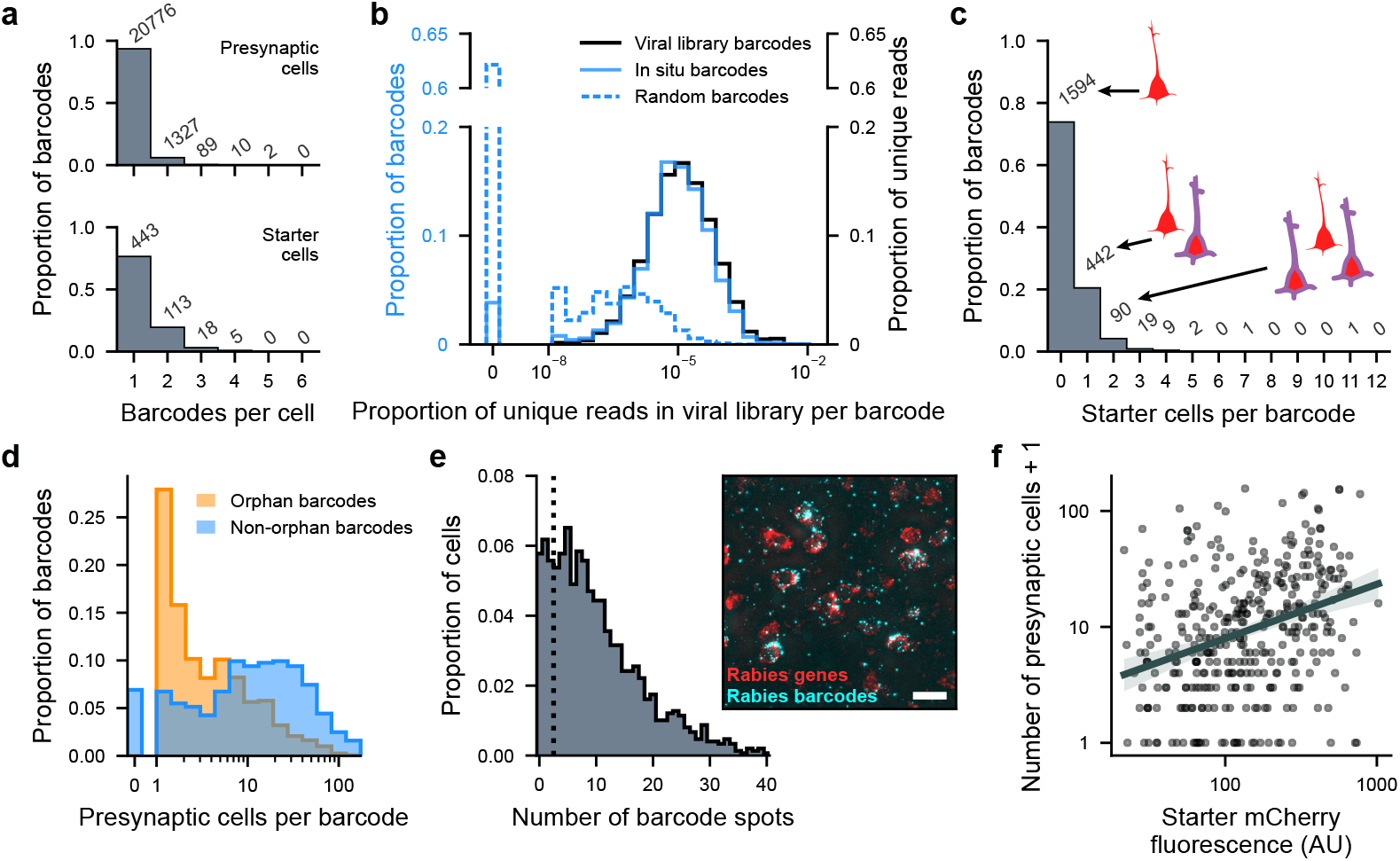
Specificity and sensitivity of *in situ* barcode detection. **a**, Number of barcodes detected in individual presynaptic and starter neurons. Most presynaptic neurons express a single barcode, while some starter neurons express >1. **b**, Barcodes detected *in situ* (solid blue line) reflect the abundance distribution of the viral library (black). In contrast, random barcodes (dashed blue line) were rarely found in the library and typically match low abundance barcodes. **c**, Number of starter cells per detected barcode sequence. **d**, Number of presynaptic cells for orphan barcodes and barcodes found in at least one starter. Orphan barcodes are typically found in a small number of cells. **e**, Number of barcode transcripts detected in rabies-infected neurons identified using probes targeting rabies viral transcripts. Dotted line indicates cells with <3 barcode spots. Inset image shows rabies viral transcripts in red and rabies barcode transcripts in cyan. Scale bar – 20 µm. **f**, mCherry fluorescence intensity in starter neurons is correlated with the efficiency of transsynaptic spread (r=0.30, p=2.6 × 10^-9^ for Pearson correlation of log-fluorescence and log (number of presynaptic cells + 1), n = 377 starters).

In total, we detected 2,158 distinct barcode sequences in rabies-infected cells. 96.2% of these barcodes precisely matched sequences in the RNA sequencing library derived from the barcoded viral prep, compared to only 37.9% of randomly generated barcodes. The probability of finding a barcode *in situ* was proportional to their abundance in the sequencing library (Fig. 3b), validating the fidelity of barcode detection *in situ*. In contrast, random barcode sequences tended to correspond to low-abundance sequences in the library (Fig. 3b).

Among these barcodes, 564 were found in starter neurons. The majority of these were found in exactly one starter cell (Fig. 3c). The remaining 1,594 orphan barcodes that were not found in any starter cells were typically found in a small number of presynaptic cells (Fig. 3d). We considered several factors, which could explain the presence of these orphan barcodes: direct infection of non starter cells during the initial virus injection, failure to detect starter neurons due to the sensitivity limits of barcode sequencing or mCherry fluorescence detection, and the death or loss of starter neurons before sequencing.

Neurons infected by the CVS-N2c strain of rabies virus have previously been shown to remain viable for at least 2 weeks (20). Since the tissue in our experiments was collected 1 week after rabies injection, extensive rabies-induced cell death is unlikely. However, we cannot exclude that some of the orphan barcodes could result from the death of starter neurons due to rabies cytotoxicity.

Orphan barcodes could also result from direct infection of mCherry-negative cells during the initial virus injection as a consequence of contamination by G-coated rabies virus or leaky TVA expression. To quantify the frequency of infection by G-coated rabies virus, we injected the barcoded rabies virus without injecting starter AAVs and quantified the number of rabies-infected neurons around the injection site (Supplementary Fig. 5a, n=2, 4 and 5 neurons). We also injected AAV FLEX-H2B-EBFP2-P2A-TVA without AAV PHP.eB hSyn-Cre or AAV FLEX-H2B-mCherry-P2A-N2cG. This was followed by rabies injection after 14 days to determine the rate of direct rabies infection due to leaky TVA expression (Supplementary Fig. 5b, n=2, 9 and 17 neurons). As the number of rabies-infected neurons detected in these experiments was far fewer than the 11,445 presynaptic cells expressing orphan barcodes observed in our *in situ* sequencing data, orphan barcodes cannot be explained by contamination from G-coated rabies virus or leaky TVA expression.

Orphan barcodes could also result from the missed detection of starter neurons due to the limited sensitivity of barcode detection. To quantify this sensitivity, we designed a panel of probes targeting the transcripts of N, P, M, and L rabies virus genes, which allowed us to reliably detect rabies-infected neurons *in situ* (Fig. 3e, inset). We found that 17.5% of rabies-infected neurons contained *<* 3 barcode spots and therefore, would not be identified in the sequencing experiments described above (Fig. 3e). Therefore, while identification of starters and presynaptic neurons may be limited by the sensitivity of barcode detection, it cannot fully account for the frequency of orphan barcodes and the fact that the majority of them are present in <5 presynaptic cells.

Alternatively, orphan barcodes could originate from starter neurons with low mCherry expression that fall below our detection threshold. Since H2B-mCherry is co-expressed in a bicistronic cassette with N2cG via a P2A ribosome skipping peptide sequence, cells with low mCherry expression are likely to express low levels of N2cG (29), resulting in inefficient transsynaptic spread (30, 31). This would explain the observation that orphan barcodes are typically found in a small number of presynaptic neurons (Fig. 3d). To explore this possibility, we selected barcodes found in exactly one starter neuron and quantified mCherry fluorescence intensity in the starter cell. We found that mCherry fluorescence was correlated with the number of presynaptic cells with the same barcode sequence (Fig. 3f). Together, these findings suggest that the excess of orphan barcodes with small numbers of presynaptic cells originates from starter neurons that fell below the mCherry detection threshold.

Among barcodes found in starter neurons, 442 were found in exactly one starter, while 122 were found in multiple starter cells (Fig. 3c). As discussed above, barcodes present in multiple starter neurons could result from initial infection by virions carrying the same sequence or transsynaptic spread between starters. To distinguish between these possibilities, we first asked whether barcodes present in multiple starter neurons were more abundant in the viral library (Supplementary Fig. 6a,b). These barcodes largely followed the abundance distribution of the viral library, with only a small excess of high-abundance sequences among barcodes present in multiple starters (6.6%, 8/122 barcodes). In addition, the median pairwise distance between starter cells sharing a barcode was shorter than between those labeled with different barcode sequences (Supplementary Fig. 6c). Together, these observations suggest that non-uniquely labeled starter cells primarily result from transmission of barcodes between starters. Barcodes present in >1 starter cell were excluded from further analyses.

### Barcoded rabies tracing reveals spatial organization of presynaptic ensembles of single neurons

We next analyzed the spatial distribution of starter and presynaptic neurons by registering *in situ* sequencing data to the Allen Common Coordinate Framework reference atlas (32). The majority (93.2%) of starter neurons were located in V1, distributed along the mediolateral axis following the locations of rabies virus injections (Fig. 4a). On the other hand, presynaptic neurons were more broadly distributed and were found in higher visual areas, retrosplenial cortex, temporal association area, auditory areas, and lateral geniculate nucleus in the thalamus. Within V1, starter neurons were most densely concentrated within layers 2/3 and 5 (Fig. 4b), potentially reflecting the tropism of AAV PHP.eB (33, 34), while presynaptic cells were more broadly distributed.

**Fig. 4.**
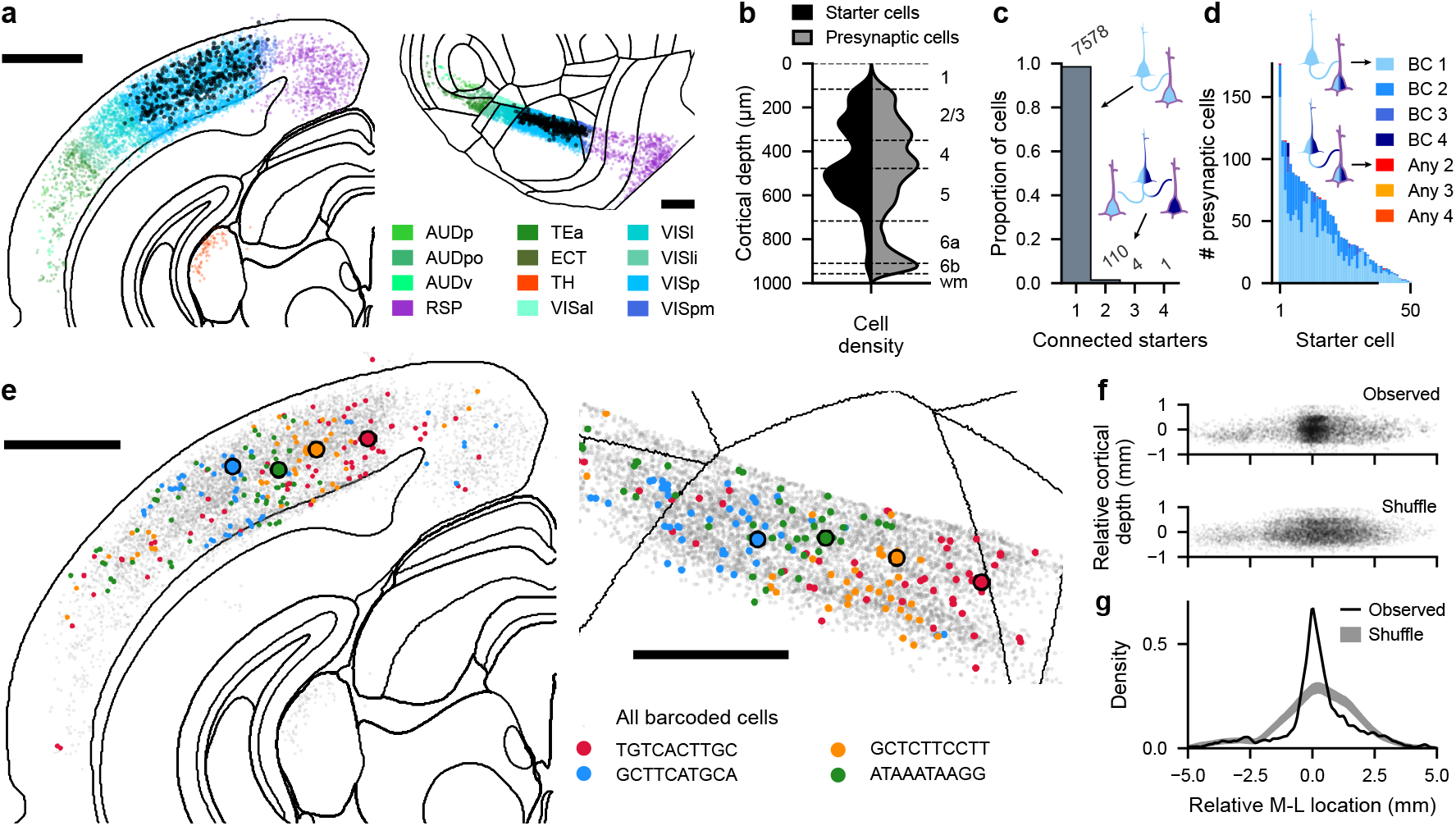
Presynaptic ensembles of barcoded starter neurons. **a**, Spatial location of starter neurons (black) and presynaptic cells. Presynaptic cells are color-coded according to brain area. Left – coronal projection, area outlines correspond to the coronal plane at the center of the sequencing volume, right – cortical flatmap. Scale bar – 1 mm. **b**, KDEs of the layer distribution of starter and presynaptic neurons in V1 in cortical flatmap coordinates. **c**, Number of starter cells connected to each presynaptic cell. **d**, Barcodes present in presynaptic cells for starter cells expressing multiple unique barcodes. Starter cells almost always only transmit one barcode to presynaptic neurons. **e**, Spatial location of starter and presynaptic neurons for 4 example barcodes. Left – coronal projection, right – cortical flatmap centered on the visual cortex. Scale bar – 1 mm. **f**, Location of presynaptic cells relative to starter neurons in cortical flatmap coordinates compared to shuffle control. **g**, KDE of relative mediolateral location of presynaptic cells with respect to starter cells. Presynaptic cells are clustered near starter cells. Shaded area indicates 95% confidence interval of the shuffle control (see Methods).

We analyzed the putative presynaptic sub-networks of starter neurons expressing barcodes that were found in exactly one starter cell. The majority of presynaptic cells were connected to one such starter neuron, although some contained barcodes from multiple starter cells (Fig. 4c). For starter neurons that expressed multiple barcodes, presynaptic neurons almost always received only one of them (Fig. 4d). This is likely a consequence of the low probability of retrograde transmission of each barcode and may also be exacerbated by superinfection exclusion after primary rabies infection has been established in a presynaptic cell (35).

We considered all neurons expressing any of the unique barcodes present in a given starter cell as a part of its presynaptic sub-network. On average, we detected 20.3 presynaptic cells for every uniquely labeled starter neuron. As the sequenced volume contained the majority of starter cells but only ∼ 44% their presynaptic inputs (Supplementary Fig. 4b), this figure underestimates the overall efficiency of transsynaptic spread.

Presynaptic neurons connected to V1 cells were found both within V1 and in other cortical areas (Fig. 4e). They tended to cluster near their starter cell (Fig. 4f-g, median distance 0.73 mm vs 1.14 mm for shuffle control, p-value = 0.002, n = 385 starter cells, 7,693 presynaptic cells, 7,814 connections, see Methods), consistent with the fact that the connection probability of cortical neurons rapidly decreases with intersomatic distance (36, 37).

### Layer- and cell-type-specific connectivity rules

Focusing on the connectivity of excitatory neurons, we analyzed the layer distribution of local (< 1 mm) presynaptic inputs as a function of the layer location of the starter cells (Fig. 5a). We first computed the input fraction, i.e., the proportion of excitatory inputs across different cortical layers for starter neurons across cortical layers, revealing layer-specific input patterns (Fig. 5b-d). Most notably, neurons in layer 2/3 were strongly recurrently connected, receiving 42% of their local excitatory inputs from other layer 2/3 cells. Their second strongest input arose from layer 4, reflecting the well-established pathway through which feed-forward sensory signals are conveyed to layer 2/3 neurons (38). Neurons in layer 5 were also recurrently connected and received strong layer 2/3 input. Finally, layer 6a received broad input from both superficial and deep layers. These findings are consistent with the canonical organization of the cortical circuit and bulk rabies tracing data (39).

**Fig. 5.**
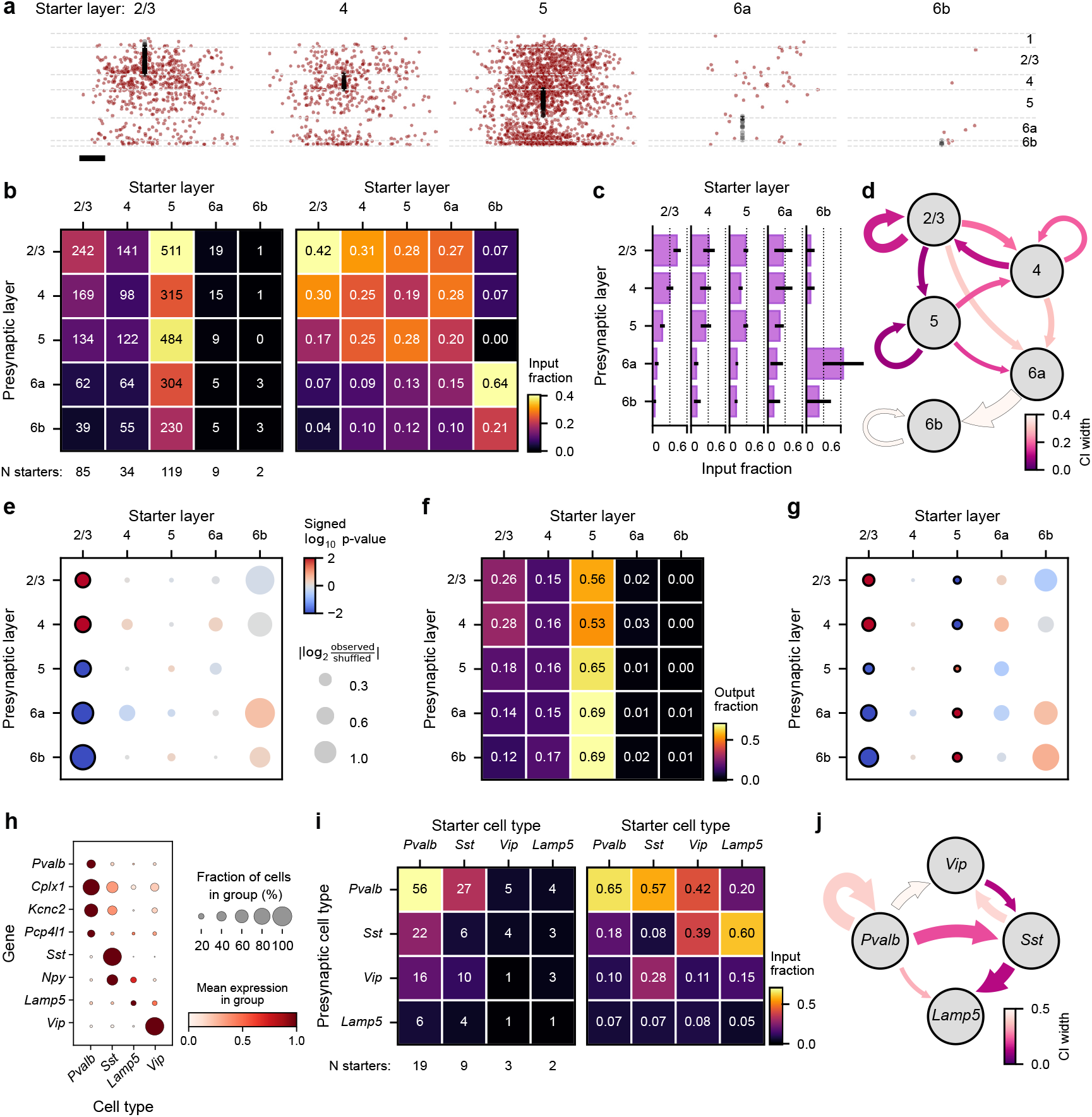
BRISC reveals layer-and cell-type-specific connectivity patterns. **a**, Relative mediolateral location and cortical depth of presynaptic cells connected to starter neurons across cortical layers in cortical flatmap coordinates. Black – starter cells; red – presynaptic cells. Scale bar – 200 µm. **b**, Number of presynaptic neurons for starter cells across cortical layers (left) and their input fractions (right). **c**, Input fractions for starter cells across cortical layers, including 95% confidence intervals computed by bootstrap resampling starter cells. **d**, Graphical representation of input fractions in **b-c**. Only connections representing >0.2 of inputs to a given starter population are shown. Arrow width is proportional to connection strength; color represents the width of the confidence interval. **e**, Input fraction compared to shuffle control. The radius of the bubbles indicates effect size, i.e., the log-ratio of observed and shuffle control input fractions. The color of the bubbles indicates significance and direction of the effect, with red and blue hues representing over- and under-representation compared to shuffle control, respectively. Solid outline indicates statistical significance (false discovery rate of 0.05). **f**, Output fraction across cortical layers. **g**, Output fraction compared to shuffle control. Notation as in panel **d. h**, Inhibitory cell type marker gene expression across *Pvalb, Sst, Lamp5*, and *Vip* interneurons. Mean normalized and log1p-transformed expression of each gene was rescaled to 1 across cell-type clusters. **i**, Number of presynaptic interneurons (left) and input fraction (right) of starter neurons across interneuron classes. **j**, Graphical representation of input fractions in **i**. Notation as in **d**.

Next, to identify over- and under-represented connections, we compared the observed connectivity matrix to a shuffle distribution, where we randomly permuted the barcodes assigned to presynaptic neurons (Fig. 5e). This analysis preserves the relative abundance of neurons across cortical layers, controlling for the tropism of starter AAVs and potential biases in the transsynaptic spread of rabies virus. It confirmed that starter neurons located in layer 2/3 preferentially received input from other layer 2/3 cells, as well as layer 4 neurons, and were less likely to receive inputs from deep cortical layers.

We next analyzed the connectivity matrix from the perspective of presynaptic neurons and computed the layer distribution of starter neurons as a function of presynaptic cell layer (Fig. 5f). As these output fractions are biased by the distribution of starter cells, we used the shuffling procedure described above, which maintains the distribution of starter cells and the number of presynaptic cells per starter, and compared the observed output fractions to the shuffle distribution (Fig. 5g). We found that layers 2/3 and 4 preferentially target layer 2/3 neurons but are less likely to target layer 5. On the other hand, layers 5, 6a, and 6b preferentially target layer 5 and are less likely to project to layers 2/3 and 4. Together with the analysis of input fractions, these results demonstrate that BRISC can identify layer-specific connectivity rules of excitatory neurons.

By sequencing the transcripts of cell-type marker genes alongside viral barcodes, BRISC makes it possible to relate input connectivity of individual neurons to their molecular identity. To demonstrate this, we focused on connectivity between GABAergic interneuron cell types in V1. We identified rabies-infected neurons belonging to 4 inhibitory neuron cell types: somatostatin-(*Sst*), parvalbumin- (*Pvalb*), vasointestinal peptide- (*Vip*) and Lysosomal Associated Membrane Protein Family Member 5 (*Lamp5*) expressing neurons (12) (Fig. 5h). We analyzed the inter-connectivity of these inhibitory cells in the same manner as for excitatory neurons by computing the input fraction onto each interneuron class (Fig. 5i-j). *Pvalb* interneurons were most frequently interconnected, consistent with connectivity measured using multiple patch clamp recordings (37). *Sst* interneurons received input from *Pvalb* and *Vip* cells, the latter connection representing a well-established disinhibitory circuit motif (37, 40). Finally, *Lamp5* and *Vip* interneurons received strong inputs from *Sst* and *Pvalb* cells. However, our sample size (3 *Vip* starter cells and 2 *Lamp5* starter cells) is insufficient to make strong conclusions.

### Long-range connectivity

Previous work has shown that the connections between the primary visual cortex and higher visual areas are topographically organized, linking areas representing the same part of the visual field (41). Our rabies injections spanned the primary visual cortex along the mediolateral axis, which largely corresponds to the azimuth axis in retinotopic space (42). Therefore, we analyzed how the spatial location of presynaptic neurons was related to the mediolateral position of their starter cell (Fig. 6a-c). Within V1, presynaptic cells tended to make connections onto starter cells at a similar mediolateral location, consistent with the spatial clustering of presynaptic cells near the starter cell. Consequently, the locations of connected presynaptic and starter cells were positively correlated within V1 (Fig. 6e). However, this relationship did not extend outside of the borders of V1. Instead, this gradient was reversed for presynaptic neurons located in cortical areas lateral to V1, where more laterally located presynaptic neurons were more likely to project to medially located starter cells. This organization reflects the reversal of the azimuth tuning gradient at the lateral boundary of V1 (Fig. 6d,f) (42, 43).

**Fig. 6.**
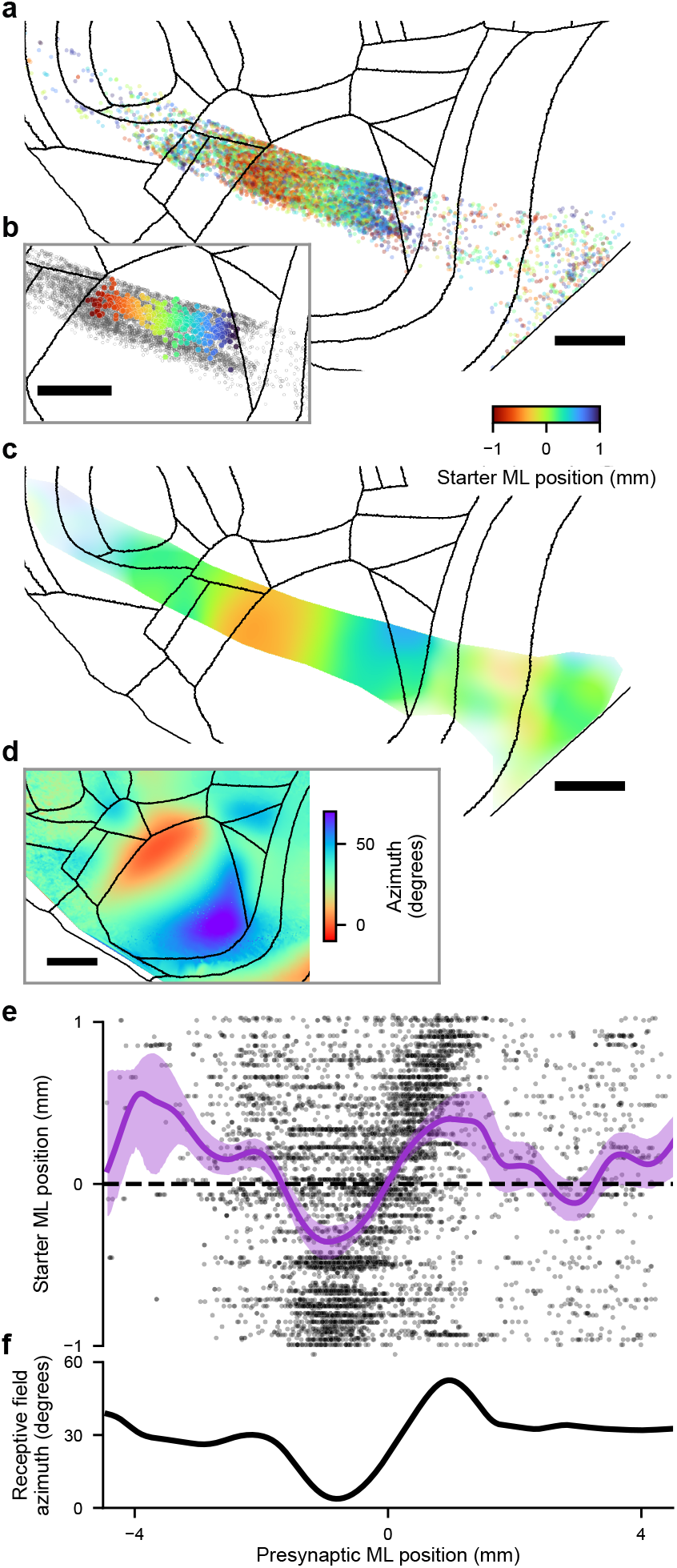
BRISC identifies topographic organization of long-range inputs onto single neurons. **a**, Flatmap projection of the location of presynaptic neurons connected to V1 starter cells, color-coded by the mediolateral (ML) position of their starter cells in flatmap coordinates. Scale bar – 1 mm. **b**, Starter cells color-coded by their ML position on the flatmap projection. Gray – all presynaptic cells. **c**, Map of average starter ML position for all cells in **a. d**, Visual azimuth map of cortical visual areas based on data from Waters et al. (44) (see Methods). **e**, Starter cell ML position as a function of the presynaptic cell position. Black – individual cells, purple – gaussian kernel density estimate. Shading indicates 95% confidence intervals computed by bootstrap resampling of starter cells. **f**, Expected preferred receptive azimuth of presynaptic cells given their spatial location (see Methods).

## Discussion

Here we describe the development of Barcoded Rabies In Situ Connectomics, BRISC, for high-throughput reconstruction of synaptic connections in the brain. Identification of individual synaptic connections using barcoded rabies viruses relies on unique labeling of starter neurons with distinct barcode sequences, which has been challenging to accomplish to date (5). To maximize the number of uniquely labeled cells, we generated high-diversity libraries of barcoded rabies viruses that could label >1,000 starter cells while ensuring that >95% of them receive distinct barcodes. The diversity of these libraries surpasses most that have been described previously (4, 5, 25). Shin et al. has recently described the generation of high complexity rabies virus libraries theoretically capable of uniquely labeling an even larger number of neurons. However, their barcode design consists of a combination of predefined index sequences, which are impractical to read out using *in situ* sequencing, limiting the potential applications of these libraries. Furthermore, sequence homology between these barcodes makes them susceptible to template switching during sequencing library preparation (45), potentially inflating the estimates of library diversity. Given the diversity of our library, the number of starter cells that can be uniquely labeled is primarily constrained by the probability of barcode transmission between them. Since this probability strongly depends on the density of starter cells (Fig. 1g), we developed a strategy to control the proportion of neurons in the target area that are infected by starter AAVs. This strategy relies on stochastic infection of starter neurons by a blood-brain-barrier permeable AAV expressing Cre recombinase, enabling us to simultaneously map the inputs across many cell types. We detected 385 uniquely labeled starter neurons, identifying 7,814 putative synaptic connections in a single animal, more than an order of magnitude higher throughput than has previously been achieved using barcoded rabies viruses (5). The resulting connectivity matrix revealed layer- and cell-type-specific connectivity rules of V1 neurons. These rules recapitulate well-established aspects of cortical connectivity, including strong recurrent intralaminar connectivity of layer 2/3 and layer 5 neurons, feed-forward connectivity across layers 4, 2/3, and 5, strong connectivity between *Pvalb*-positive interneurons, and inhibition of *Sst* interneurons by *Vip*-positive cells.

By densely sequencing the sections containing the majority of starter neurons, we could identify and eliminate barcodes that were present in more than one starter cell. However, some of the barcodes detected in exactly one starter neuron may have been present in other starter cells that escaped detection due to limitations of sensitivity of *in situ* sequencing or low mCherry expression. Our results suggest that low mCherry expression is associated with low efficiency of barcode transmission, presumably due to low expression of the G glycoprotein. As starter cells in this category are unlikely to transmit their barcodes to a significant number of presynaptic cells, they are unlikely to substantially impact the analysis of BRISC data.

The efficiency of BRISC is challenging to estimate due to the lack of ground truth measurements of the number of distinct presynaptic partners of pyramidal neurons. Mouse visual cortex neurons are estimated to have ∼ 2,000 dendritic spines (46). However, connected pairs of neurons often make multiple synaptic contacts, with light microscopy reconstructions estimating that connected pyramidal neurons on average share ∼ 5 synapses (47). Together, these figures suggest that visual cortex cells receive synaptic inputs from ∼ 400 distinct presynaptic neurons, although this estimate does not include inhibitory connections. As barcoded rabies tracing labels ∼ 50 presynaptic neurons per starter cell, this likely constitutes ∼ 10% of their presynaptic partners. Our results suggest that increasing the levels of G glycoprotein expression would increase the sensitivity of starter cell detection as well as the number of presynaptic neurons labeled by each starter, enhancing the throughput of BRISC and providing a more comprehensive sample of inputs onto each neuron.

By simultaneously reading out connectivity and gene expression patterns of individual neurons, BRISC has the potential to comprehensively characterize cell-type-specific wiring rules. In addition, BRISC makes it possible to correlate gene expression and connectivity at the single-cell level, potentially identifying connectivity rules that do not align with established transcriptomic cell types. The scalability of BRISC enables the application of connectomics to understand neural mechanisms of neurological or psychiatric diseases by analyzing wiring patterns in disease models to identify defects in cell-type-specific wiring rules.

As BRISC preserves spatial information, it can be integrated with other imaging modalities, including *in vivo* multi-photon imaging. By registering *in vivo* imaging volumes to *in situ* sequencing cryosections (28), BRISC could be applied to simultaneously determine functional properties and input connectivity of individual cells at scale, providing ground-truth data to constrain circuit models of cortical computations.

BRISC also enables the systematic comparison of connectivity rules across multiple brain areas or experimental subjects. Since BRISC depends entirely on viral vectors and not transgenic animals, it is straightforward to apply to other brain areas and potentially other animal species. This creates new opportunities for the use of comparative connectomics to systematically explore how microcircuit organization differs across the brain and the evolutionary tree.

## Author contributions

P.Z. and A.Be. conceived the study. A.Be. and M.S. generated barcoded rabies libraries. A.Be. and A.Bl. carried out barcoded rabies tracing experiments. A.Be, A.Bl., and P.Z. analyzed the data and wrote the manuscript.

## Acknowledgments

This work was supported by the Francis Crick Institute which receives its core funding from Cancer Research UK (CC2108), the UK Medical Research Council (CC2108), and the Wellcome Trust (CC2108). We would like to thank Sophie Wood for performing some of the surgical procedures, Scott Lighterness for tail vein injections, Matt Renshaw for help in designing the microscope, Andreas Schaefer, Johannes Kohl, Dinu Florin Albeanu, and James Briscoe for comments on the manuscript, Andrew Murray for providing the CVS-N2c-ΔG-mCherry rabies genomic plasmid that was used as the backbone for the barcoded rabies viral libraries (Addgene plasmid # 73464). We thank the staff of the Making Lab, Genomics, Light Microscopy, and Biological Research Facilities at the Francis Crick Institute for supporting this work. This preprint was typeset in LaTeX based on [ a template created by Ricardo Henriques](https://www.overleaf.com/latex/templates/henriqueslab-biorxiv-template/nyprsybwffws).

## Data and code availability

Preprocessed data required to reproduce the analyses will be deposited and made publicly available prior to publication. Both preprocessed and raw data are available from Petr Znamenskiy upon request. Original code required to reproduce the analyses is available on Github at https://github.com/znamlab/brisc. Code for processing raw *in situ* sequencing data is available at https://github.com/znamlab/iss-preprocess and https://github.com/znamlab/iss-analysis.

## Methods

### Animals

All animal procedures were conducted in compliance with the guidance and regulations set forth by the UK Home Office according to the Animals Scientific Procedures Act 1986 (PPL PP4882546) and approved by the Animal Welfare Ethical Review Body at the Francis Crick Institute. Male and female C57Bl/6 mice aged between 12 and 15 weeks were used for barcoded rabies experiments. Ai14-tdTomato Cre-responder mice (B6;129S6-Gt(ROSA)26Sor^tm14(CAG-tdTomato)Hze/J^, JAX #007908) were used for AAV-PHP.eB-Cre titration experiments.

### Barcoded rabies viral library production

#### Plasmid library production

A CVS-N2c-ΔG-mCherry plasmid (Addgene plasmid # 73464, a gift from Dr Andrew Murray) was used as the basis for producing barcoded rabies libraries. The extended 3’ sequence that is necessary to generate the super-cutter variant of the hepatitis delta virus antigenomic ribozyme (HDVagrz) was introduced into the plasmid using a Q5 site-directed mutagenesis kit (New England Biolabs, E0554S). An oligonucleotide was inserted containing a sequence complementary to the 5’ region of the hepatitis delta virus antigenomic ribozyme, along with 5 extra bases (ACGGA).

Barcodes were then inserted into the modified CVS-N2c-ΔG-mCherry plasmid downstream of the mCherry transgene by inverse PCR (4) using a pair of forward and reverse primers (Bipartite_barcode_F and Bipartite_barcode_R for RV2, Barcode_F and Barcode_R for RV35, Supplementary Table 1) comprising a 5’ tail of six protective bases flanking an EagI restriction site, a genome complementary 3’ sequence, and random nucleotides forming the barcode (10 bases in both primers for RV2, 20 bases in the forward primer only for RV35). These random nucleotides were incorporated using IDT’s “hand-mixed” option at a final ratio of N(25:25:25:25) to account for differences in coupling rates of each nucleotide. To produce large amounts of linear barcoded plasmid DNA, PCR was performed in a 96-well PCR plate with 25 µl per well. This mix was comprised of 1,250 µl Q5 Hot Start High-Fidelity 2X Master Mix, 125 µl forward primer (10 µM), 125 µl reverse primer (10 µM), 500 µl template plasmid (supercutter CVS-N2c-ΔG-mCherry, 0.4 ng/µl) and 500 µl molecular grade water. The following thermocycling conditions were used to generate the linear plasmid constructs with EagI 5’ and 3’ ends: 98 °C - 30 s, 35 cycles of 98 °C for 10 s, 72 °C for 30 s, 72 °C for 450 s, followed by 72 °C for 2 mins and 4 °C hold.

To size-select the resultant linear products, the PCR reaction was loaded on an agarose gel, and gel extraction was performed with a Wizard SV Gel and PCR Clean-Up kit (Promega, A9281). Secondary purification was then carried out with DNA Clean & Concentrator-5 columns (Zymo Research, D4004). Typically, around 20 µg of linear DNA was recovered from a single 96-well plate.

The purified PCR product was then digested at 37 °C for 1 hour to expose sticky ends for ligation with the following reaction mix: 0.5 µl EagI-HF (New England Biolabs, R3505L), 0.25 µl DpnI (New England Biolabs, R0176L), 500 ng linear barcoded DNA, 5 µl CutSmart buffer (New England Biolabs, B6004S) and molecular-grade water up to 50 µl. Heat inactivation was then carried out at 80 °C for 20 minutes. The sticky-ended products were then ligated into open-circular plasmids by adding T4 ligase (New England Biolabs, M0202T) and ATP (Thermo Fisher Scientific, PV3227) to the reaction mix at a ratio of 0.1 µl ligase and 5 µl ATP per 44.9 µl of sticky-ended product and incubating at 4 °C for 2 hours, followed by heat inactivation at 65 °C for 20 minutes. Residual linear DNA was then digested by adding Exonuclease V (New England Biolabs, M0345S) to the reaction mix (1 µl ExoV, 1.4 µl NEBuffer4 (New England Biolabs, B7004S) and 6.4 µl ATP) and incubating at 37 °C for 1 hour, followed by heat inactivation at 70 °C for 30 minutes. The resulting product was purified using DNA Clean & Concentrator-5 columns.

Barcoded open circular plasmids were transformed into MegaX DH10B T1R Electrocompetent cells (Thermo Fisher Scientific, C640003) to amplify and supercoil them for mammalian transfection. 40 µl of electrocompetent cells were mixed with 400 ng of circularized DNA per cuvette, added to pre-chilled electroporation cuvettes on ice (BIO-RAD, 1652082), and immediately electroporated at 2.0 kV, 200 Ω, 25 µF using a GenePulser XCell (BIO-RAD, 1652662). 1 ml of pre-warmed SOC media was then added to each cuvette and placed in a round-bottomed tube in a shaking incubator for 1 hour at 37 °C. All samples were then pooled, diluted to 20 ml with LB broth, and then 2 ml was spread evenly across each of 10 prewarmed LB Carbenicillin (100 µg/ml) 245 mm Square BioAssay plates (Corning, 431111). The plates were incubated overnight at 30 °C, and then all colonies were scraped from the plates. The bacteria were centrifuged to collect a pellet. ZymoPURE II maxiprep kits (Zymo Research, D4202) were used to extract endotoxin-free plasmid DNA.

### Rabies packaging and pseudotyping

Initial viral rescue attempts resulted in the RV2 library (Supplementary Fig. 1a,b). Rescue from barcoded plasmids was carried out in 2 x 10 cm plates of HEK293T cells transfected with 7 µg barcoded rabies genome plasmid, 6 µg pCAG-T7-polymerase, 12 µg pCAG-N2c N, 6 µg pCAG-N2c P, 6 µg pCAG-N2c L, and 6 µg pCAG-N2c G plasmids in 1000 µl of Opti-MEM with 130 µl PEI MAX (1 mg/ml), mixed and incubated for 20 minutes before adding to the cells. The following day, the media was exchanged with fresh DMEM / 10% FBS. At 7 days post-transfection, cells were split in DMEM / 10% FBS at a ratio of 1:3.3 into 2 x 10 cm plates and 2 x 15 cm plates. At 8 days post-transfection, the cells were re-transfected with 40 µg pCAG-N2c-G in 2000 µl of Opti-MEM and 120 µl PEI MAX (1 mg/ml). This was added to the cells and incubated for 6-8 hours before replacing with fresh DMEM / 10% FBS media. At 17 days post-transfection, the supernatant from the HEK293T plates was transferred to 2 x 15 cm dishes of BHK-EnvA cells, seeded at a density of 1 x 10^7^ cells per dish for pseudotyping. Cells were trypsinized and replated with fresh media after 6 hours to remove residual G-coated virus. At 21 days post-transfection, the supernatant was collected, filtered through a 0.45 µm filter, and centrifuged at 50,000 g for 2 hours at 4 °C. Pellets were resuspended in 300 µl PBS, and the viral titer was determined by applying serial dilutions of rabies virus to HEK293T-EnvA (48) and HEK293T cells, counting mCherry+ cells.

We then optimized the rescue protocol to increase library diversity, generating library RV35. Specifically, we used a stable cell line expressing the N2c-glycoprotein (HEK293TG) to perform initial viral rescue and decreased the duration of viral amplification steps to reduce clonal amplification biases. To generate HEK293TG cells, HEK293T cells were transduced with a 3rd generation EF1a-N2cG-puromycin lentiviral vector containing the rabies glycoprotein gene flanked by long terminal repeat sequences for integration into the genome. Lentiviruses were produced by mixing a pEF1a-N2cG-IRES-Puro transfer plasmid (8.3 µg) with packaging and envelope plasmids (pLP1-9.6 µg, pLP2-9.0 µg, VSV-G-6.4 µg) to transfect 2 × 10^7^ HEK293T cells in a 15-cm culture dish with 100 µl Lipofectamine 2000 (Invitrogen, 11668019). Viral supernatants were collected and filtered at 2 and 3 days post-transfection, before being pooled and concentrated with a solution of 10% w/v PEG-8,000 (Merck, 81268) and NaCl (88 mM). Aliquots of lentivirus particles were titered by ddPCR (49). HEK293T cells were then transduced with lentivirus at an MOI of 0.3. After 48 hours, cells were maintained in media containing 2 µg/ml of puromycin (Sigma, P9620) until single colonies were isolated to select for cells that stably expressed the glycoprotein transgene. This was confirmed by infecting clonal colonies with glycoprotein-coated CVS-N2c-(ΔG)-mCherry rabies virus at low MOI. After several days, supernatant was taken from the wells and added to fresh HEK293T cells. Observation of mCherry fluorescence in the HEK293T cells was used to confirm successful packaging of glycoprotein-coated rabies virus.

HEK293TG cells were transfected with the barcoded rabies genome plasmid along with plasmids containing the rabies genes N, P, L and G, as well as T7 polymerase. Six-well dishes were first seeded with HEK293TG cells at a density of 6 x 10^5^ cells/well in DMEM / 5% fetal bovine serum (FBS). The transfection mix was prepared containing 6 µg pTIT-N, 3 µg pTIT-L, 5 µg pTIT-P, 6 µg pTIT-G, 8.5 µg pCAGGS T7 and 12 µg barcoded plasmid library in 430 µl of Opti-MEM. Then, 85 µl of PEI MAX (1 mg/ml) was added to the mix, incubated at room temperature for 20 minutes, and 80 µl of this was added to each well. The following day, the media was exchanged with fresh DMEM / 10% FBS. Three days after transfection, the cells were split in DMEM / 10% FBS at a ratio of four wells to a T175 flask. When cells reached 70% confluency (6-7 days post-transfection), they were transfected with the glycoprotein plasmid in the following transfection mix: 13 µg CMV-RG, 1 ml Opti-MEM, 40 µl of PEI MAX (1 mg/ml). This was added to the cells and incubated for 6-8 hours before replacing with fresh DMEM / 10% FBS media. Supernatant was collected 9-10 days after the initial transfection and filtered through 0.45 µm filters. The supernatant was titered by serial dilution onto HEK293TG cells in 6-well plates, with a typical titer of 1 x 10^7^.

This supernatant was then used to infect BHK cells stably expressing the EnvA coat protein (BHK-EnvA-GFP, (48)) for pseudotyping. BHK-EnvA cells were seeded in 150 mm dishes at a density of 1 x 10^7^ cells per dish in DMEM / 10% FBS. Supernatant containing barcoded G-coated rabies virus was added to each dish at an MOI of 0.3. The cells were incubated with supernatant for 6-8 hours and then washed with PBS four times, then split and replated in DMEM / 5% FBS at a ratio of 1:3. Supernatant containing EnvA-pseudotyped virus was collected and filtered through 0.45 µm filters 2 days after infection and fresh DMEM / 10% FBS was added to the plates. The supernatant was centrifuged at 70,000 g in an SW32.1 rotor for 2 hours at 4 °C and the viral pellets were resuspended in approximately 250 µl PBS. This was repeated 3 days after infection. The titer of the concentrated viral prep was determined by applying serial dilutions of the virus to HEK293T-EnvA (48) and HEK293T cells to quantify pseudotyped and glycoprotein-coated viral titers, respectively.

#### Barcoded library sequencing

The diversity of the barcoded plasmid and viral libraries was quantified using a unique molecular identifier (UMI) tagging approach. For viral libraries, viral genomic RNA was extracted using the Quick DNA/RNA Viral Kit (Zymo Research, D7020) and eluted into 15 µl of RNase-free water. Purified viral barcode RNA was then reverse transcribed using Superscript IV reverse transcriptase (Thermo Fisher Scientific, 18090050) at 50 °C for 10 minutes with 1 µl of 2 µM UMI_P5_Truseq primer (Supplementary Table 1), before heat inactivation at 80 °C for 10 minutes. RNA was subsequently digested with RNase H (New England Biolabs, M0297S) at 37 °C for 20 minutes. The primer used for reverse transcription (UMI_P5_Truseq, Supplementary Table 1) contained a 5’ 22 bp region of the P5 Truseq sequence for nested adaptor PCR, a 12 bp random UMI sequence and 20 bp of rabies homologous sequence for binding upstream of the barcode.

The same UMI-P5-Truseq primer was used to produce UMI-tagged amplicons from the barcoded plasmid libraries in a two-cycle limited PCR reaction. Barcoded plasmid (5 ng) was added to 16 µl Q5 Hot Start High-Fidelity 2X Master Mix (New England Biolabs, M0494S) with 1.25 µl each of UMI_P5_Truseq and P7_universal_reverse primers (Supplementary Table 1) and PCR thermocycled with the following conditions: 98 °C - 3 mins, two cycles of 98 °C - 30 s, 67 °C - 20 s, 72 °C - 30 s, followed by 72 °C - 2 mins and 12 °C hold.

The resultant cDNA from the viral samples or amplicons from barcoded plasmids was size-selected and purified using KAPA Pure Beads (Roche, KK8000) in a 1:1 ratio, eluting in 25 µl of RNase-free water. To determine the optimal cycle number for library amplification, the amplification curve inflection point was determined for each sample by qPCR. 2.5 µl of each sample was added to a master mix of 1 µl 2.5X Thiazole Green (Biotium, 40086), 1 µl 2.5X ROX reference dye (Thermo Fisher Scientific, 12223-012), 12.5 µl Q5 Hot Start High-Fidelity 2X Master Mix (New England Biolabs, M0494S), 1.25 µl 10 µM IDT_i7_1 primer (Supplementary Table 1) for plasmid amplicons or P7_universal_reverse for viral single-stranded cDNA, 1.25 µl 10 µM IDT_i5_1 primer and 5.5 µl of molecular grade water. The following thermocycler conditions were used: 98 °C - 3 mins, followed by 40 cycles of 98 °C - 10 s, 68 °C - 20 s, 72 °C - 30 s.

Subsequently, the remaining 22.5 µl of each sample was added to a mix of 25 µl Q5 Hot Start High-Fidelity 2X Master Mix, 2.5 µl 10 µM IDT_i7_1-10 primer for plasmid amplicons/P7_universal_reverse for viral single-stranded cDNA, and 2.5 µl 10 µM IDT_i5_1-10 primer. The IDT_i5_X and IDT_i7_X primers each contain 8 bp index sequences derived from the rhAmpSeq index primer set (Integrated DNA Technologies) for dual-index demultiplexing alongside the Illumina P5 and P7 adapter sequences, respectively. Thermocycling conditions for this Illumina adaptor PCR were: 98 °C - 3 mins followed by cycles of 98 °C - 10 s, 68 °C - 20 s, 72 °C - 30 s, then 72 °C - 2 mins. The number of cycles was equal to the qPCR inflection point, less 3 cycles to account for the 10X (∼2^3^) increase in target concentration.

For viral amplicons, the P7-indexed adaptor sequences were added in a final PCR step. The UMI-tagged viral amplicons were first gel-purified with a Wizard SV Gel and PCR Clean-Up kit (Promega, A9281) and eluted with 20 µl water. Then, the same PCR mix was used as described above for the barcoded plasmid index PCR, adding the IDT_i7_X primer and the matching IDT_i5_X for each sample. Thermocycling conditions were likewise the same as the Illumina adaptor PCR, but with a fixed number of 3 cycles. The indexed viral and plasmid amplicons were then gel-purified with a Wizard SV Gel and PCR Clean-Up kit (Promega, A9281) and eluted with 25 µl water. The concentration and size distribution of these amplicon libraries were quantified with a High Sensitivity D1000 ScreenTape kit (Agilent, 5067-5584) on an Agilent 2200 TapeStation system. Samples passing QC were sequenced on an Illumina NovaSeq 6000/NovaSeq X using Illumina TruSeq primers for paired-end 100 bp sequencing at a depth of 10-50 million reads per sample.

#### Quantification of barcoded library diversity

To analyze sequenced libraries, low-quality reads were first removed from raw demultiplexed .fastq files with Trimmomatic 0.36, removing reads with an average PHRED score per base of less than 20 in a 6-base sliding window and removing any reads with a length of less than 101 bp. The sequences flanking the barcode site were then matched to their expected sequences, and any mismatched sequences were discarded to remove frameshifts. Next, the UMI sequences were used to de-duplicate the remaining barcode sequence reads to account for PCR amplification bias. The remaining sequences represented individual species of barcode RNA/DNA found in the viral or plasmid samples. To account for sequencing errors and PCR-induced mutations that may be present in some of these sequences, closely related barcodes were collapsed into their most abundant “parent” sequences using a method adapted from Kebschull et al. (50). First, barcode sequences were aligned to all other barcodes in the library with a maximum hamming distance of 2 using Bowtie2 (2.5.1-GCC-12.3.0, (51)). Related sequences were then collapsed onto the barcode with the highest UMI count, summing together all of the counts. Previously published barcode library data (Supplementary Fig. 1a,b) was acquired from deposited data sources: GSM4519333 (25), Source Data 1f 170330_pSPBN_GFP_B19EnvA (4), SRR23310757 (5), and GSM8524116/ GSM8524119 (7).

### Surgical procedures

For barcoded rabies tracing experiments, Metacam (33 mg/l) was provided via drinking water before and for 3 days following surgery as peri- and post-operative analgesia. Prior to anesthesia, 50 µl AAV-PHP.eB-Syn-Cre (4.5 x 10^13^ vg/ml) diluted 1:330 in filter-sterilized PBS was administered via tail vein injection. Anesthesia was then induced and maintained with 1.5–3% isoflurane, and mice received subcutaneous injections of buprenorphine (0.1 mg/kg). Mice were then secured in a stereotactic frame, and a small craniotomy was created at the site of the injection. Virus injections were performed using a Nanoject III injector at rates not exceeding 14 nl/min. Helper viruses FLEX-H2B-mCherry-P2A-N2cG (3.1 x 10^13^ vg/ml) and AAV1-FLEX-H2B-EBFP2-P2A-TVA (1.8 x 10^13^ vg/ml), diluted in sterile PBS to 1:20 and 1:100 respectively, were delivered at three injection sites at −3.5 mm from Bregma and 3.0, 2.4 and 1.8 mm from midline in the left hemisphere, 300 and 500 µm below brain surface (60 nl per depth). Thirteen days later, barcoded CVS-N2c-mCherry (6.0 x 10^8^ IFU/ml) was injected at the same coordinates (500 nl per depth). Seven days later, mice were euthanized via intraperitoneal injection of sodium pentobarbital (600 mg/kg). Brains were extracted and rapidly frozen in OCT compound-filled cryomolds using a bath of absolute ethanol and dry ice, and stored at −80°C.

TVA leak and rabies glycoprotein-coated contamination control experiments (Supplementary Fig. 5) followed the above protocol but excluded tail vein injections and omitted either FLEX-H2B-mCherry-P2A-N2cG or both helper viruses, respectively. Seven days after rabies injection, mice were terminally anesthetized with sodium pentobarbital (600 mg/kg) and then perfused with ice-cold PBS, followed by 4% paraformaldehyde in PBS. Brains were then post-fixed overnight in 4% paraformaldehyde before imaging.

For AAV-PHP.eB-Cre titration experiments (Supplementary Fig. 1c), Ai14 mice (B6;129S6-Gt(ROSA)26Sor^tm14(CAG-tdTomato)Hze/J^, JAX #007908) received 50 µl tail vein injections of AAV-PHP.eB-Syn-Cre (4.5 x 10^13^ vg/ml) at 1:100, 1:330, 1:1,000, or 1:3,300 dilution in sterile PBS. Two weeks after tail vein injections, mice were terminally anesthetized with sodium pentobarbital, perfused and fixed as above.

For quantification of barcoded rabies spread using serial two-photon tomography (Fig. 1h-j), the procedure for barcoded rabies sequencing surgeries was also performed as described above with the following differences: a single local injection of AAV1-hSyn-Cre (7.6 x 10^13^ vg/ml) diluted 1:5000 in sterile PBS was combined with the helper viruses diluted as above and injected at coordinates of 3.5 mm AP, 2.5 mm ML, 300 and 500 µm below brain surface (60 nl per depth). Seven days later, a pair of 500 nl injections of barcoded RV2 CVS-N2c-mCherry (2.8 x 10^7^ IFU/ml) were administered at the same stereotactic coordinates. After seven more days, mice were euthanized, perfused and fixed as detailed above.

For quantification of the spatial distribution of starter neurons in intracerbral/intravenous injections of AAV-Cre (Fig. 1k-m), the procedure for barcoded rabies sequencing surgeries was also performed as described above with the following differences: wild-type mice received either a 50 µl tail vein injection of AAV-PHP.eB-Syn-Cre (4.5 x 10^13^ vg/ml) at 1:330 dilution in sterile PBS (intravenous), or a single local injection of AAV1-hSyn-Cre (7.6 x 10^13^ vg/ml) diluted 1:5000 in sterile PBS (intracerebral) at coordinates of 3.8 mm AP, 2.4 mm ML 300 and 500 µm below brain surface (60 nl per depth). For both conditions, AAV FLEX- H2B-mCherry-P2A-N2cG (3.1 x 10^13^ vg/ml), diluted in sterile PBS to 1:20 was also injected at the same location on the same day. After 14 more days, mice were euthanized, perfused and fixed as detailed above.

### Serial section two-photon tomography

To quantify the spread of barcoded rabies virus and determine the optimal parameters for AAV1-hSyn-Cre injections, PFA-fixed brains were imaged using a serial sectioning two-photon microscope (52, 53). Sections were imaged in a bath of 50 mM pH 7.4 phosphate buffer. The microscope was controlled by ScanImage Basic (MBF Bioscience), using the custom software wrapper BakingTray to control imaging parameters (54). Image tiles were assembled by StitchIt (55). To quantify AAV PHP.eB Cre-positive cells, images were acquired at an XY resolution of 1 or 2 µm with optical sectioning intervals of 5 or 8 µm and vibratome sections at 40 µm intervals. Cell detection throughout the brain was performed either using Cellfinder (56) (Fig. 1h,j, Supplementary Fig. 4b) or manually using napari (Fig. 1l,m, Supplementary Fig. 5).

As nuclear fluorescence of starter cells could not be detected from the serial section two-photon tomography dataset directly, the slices were collected, counterstained with DAPI and imaged on a confocal microscope (LSM 880, Zeiss) with voxels of 0.2 x 0.2 x 1.6 µm. Starter cells were manually identified from their brighter nuclear H2B-mCherry signal compared to the cytoplasmic rabies mCherry expression. To display both the high-intensity nuclear staining and fainter cytoplasmic fluorescence (Fig. 1i), RGB images were formed by collating the DAPI channel (in blue) with the mCherry data duplicated in both the red and green channels with different contrast limits (1,000 and 10,000 gray values, respectively).

### *In situ* sequencing

#### Primer and padlock design

Cell type marker genes were selected using SMART-Seq2 RNA expression data from neurons in the mouse visual cortex (12) as a reference. Candidate genes were selected by first filtering to remove genes with low mean expression. Then, a modified version of a greedy combinatorial search algorithm (57) was used to iteratively select the top 80 genes that maximized classification accuracy across the reference cell type clusters. Additional sixteen genes were then manually added to the genes selected by the greedy algorithm, including established visual cortex cell type marker genes (*Chodl, Cux2, Fezf2, Foxp2, Rorb, Vip*), additional marker genes that are differentially expressed in cell types where the canonical marker is downregulated by rabies infection (58) (L5 NP - *Tshz2*, Sst - *Grin3a* / *Reln*), and additional markers to improve the classification of neurons within clusters (L2/3 IT neurons - *Rrad*), or between subclasses (L5 IT/PT/NP - *Etv1, Tle4*, L6 IT/CT - *Rasl10a, Hs3st4*, Vip/Sst - *Grik1, Sorcs3*).

Multiple padlock probes were designed for each target gene (Supplementary Table 3). The number of probes designed for a given gene was calculated based on the expression level of each gene relative to *Sst* in a reference V1 SMARTseq scRNAseq dataset (12):

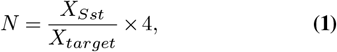

where *N* is the number of padlock probes, *X*_*Sst*_ is the mean expression of the *Sst* gene from all cells in the *Sst* cell type subclass, and *X*_*target*_ is the mean expression of the target gene in its highest expressing cluster in the reference scRNAseq dataset. This was limited to an upper bound of 12 padlocks per gene and a lower bound of 4 padlocks per gene. Padlock sequences were generated using a modified version of the [Multi padlock design](https://github.com/Moldia/multi_padlock_design) tool (57). Briefly, for each target gene, the tool determined the consensus sequence from all transcript variants in the [mouse RefSeq transcriptome](https://ftp.ncbi.nlm.nih.gov/refseq/M_musculus/mRNA_Prot/) and tested every possible binding sequence in a sliding window along the transcript. Candidate binding sequences consist of two adjacent padlock arm regions (20 bp per arm). The arms of each candidate sequence were filtered to be within a defined melting temperature range (60 - 75°C after formamide-containing hybridization buffer correction, melting temperature calculated assuming 0.1 µM probe, 0.075 M Na^+^, 0.01 M Mg^2+^ and 20% formamide). The specificity of each candidate sequence was then tested with a BLAST search against the whole mouse transcriptome to ensure padlocks do not exhibit non-specific binding. A 4-base pair minimum distance between padlock candidates was also specified to avoid steric hindrance of bound padlocks.

All padlocks had a conserved backbone sequence containing a shared sequencing primer binding site directly upstream of a 7-base gene barcode, unique to each gene. Gene barcodes were selected from a pool of previously designed “GII codes” (27) with a minimum hamming distance of 2 between each barcode. Padlock probes for the highly expressed genes *Slc17a7, Gad1, Sst* and *Vip* contained unique binding sites in place of the sequencing primer site to allow for direct readout of these genes with fluorophore-conjugated hybridization probes separately from the sequencing rounds of the other genes. Padlocks were obtained from IDT as 4 nM ultramer sequences with 5’ phosphorylation.

A reverse transcription primer was designed for each padlock by taking the reverse complement of the 5’ arm of each padlock (Supplementary Table 3). Reverse transcription primers were obtained from IDT as standard desalted oligonucleotides.

#### Brain cryosectioning

Serial sections were cut at 20 µm thickness with MX35 Ultra disposable microtome blades (Epredia, 3053835) at −13 °C specimen temperature / −15 °C chamber temperature using a Bright OTF5000 cryostat (Bright Instruments, R080) and mounted onto poly-l-lysine coated 25 x 75 mm #1.5H coverslips. Ten sections were mounted per coverslip and then stored at −80 °C.

#### Flowcell assembly and tissue preparation

Flowcells were assembled from 26 x 76 mm microscope slides with pre-drilled 1 mm holes placed at each end for fluid flow. Ports were cast from polydimethylsiloxane, punched with 1.5 mm holes and plasma bonded over the holes drilled in the microscope slides. A laser-cut 140 µm-thick adhesive gasket (Grace Biolabs) was mounted on the other side of the microscope slide.

The protocol for *in situ* reverse transcription and amplification of transcripts from tissue was adapted, combining elements of BARseq2 (27) and coppaFISH (28). Coverslips with cryosections were removed from 80 °C and immediately immersed in 4% PFA in PBS for 30 minutes. They were then immersed in PBS to wash and then washed twice in PBST (PBS with 0.5% Tween20 (Sigma Aldrich, P1379)). Next, they were immersed in a series of ethanol solutions, incubating for 5 minutes each in 70%, 85% and 100% ethanol at room temperature. The coverslips were transferred to fresh Falcon tubes containing 100% ethanol and stored overnight at 4°C. The following day, the coverslips were removed from the ethanol, and once dried, each flowcell was assembled by pressing the coverslip onto the adhesive gasket backed with the microscope slide to form an enclosed chamber. Assembled flowcells had a volume of 180 µl and were washed with PBST three times to rehydrate the tissue and remove any bubbles from within the flowcell.

#### Reverse transcription

A reverse transcription mix was then added to each flowcell. The mix contained pooled gene-specific reverse transcription primers (60 nM per primer), 1 µM rabies barcode reverse transcription primer (RTAB0067, Supplementary Table 3), 1 U/µl RiboLock RNase Inhibitor (Thermo Fisher Scientific, EO0381), 0.2 µg/µl non-acetylated BSA (Thermo Fisher Scientific, AM2616), 0.5 mM dNTP (Thermo Fisher Scientific, R0192), 20 U/µl SuperScript IV reverse transcriptase (Thermo Fisher Scientific, 18090200), 5 mM DTT (Thermo Scientific, 707265) in 1X SuperScript IV buffer. Samples were incubated at 45°C overnight in a humidified chamber to reduce evaporation.

#### Padlock ligation

The following day, samples were washed in PBST once and the cDNA from reverse transcription was crosslinked by incubating the samples in 4% PFA in PBS for 30 minutes at room temperature. Samples were then washed in PBST twice to remove excess PFA. RNA template digestion, padlock hybridization to cDNA and padlock ligation were carried out in the same reaction. First, a gene padlock ligation mix was added to the samples, containing pooled gene padlocks (60 nM of each padlock, Supplementary Table 3), 0.4U/µl RNaseH (New England Biolabs, M0297L), 20% Formamide, 50 mM KCl, 0.5U/µl Ampligase (Lucigen, A0102K) in 1X Ampligase buffer. Samples were incubated for 40 minutes at 37°C, followed by 1 hour at 45°C. Then, the gap-filling/ligation mix containing 1 µM rabies barcode padlock (OAB0973), 0.4U/µl RNaseH (New England Biolabs, M0297L), 20% Formamide, 50 mM KCl, 0.25 µg/µl non-acetylated BSA (Thermo Fisher Scientific, AM2616), 50 µM dNTP (Thermo Fisher Scientific, R0192), 5% glycerol, 0.001U/µl Phusion polymerase (Thermo Fisher Scientific, F530L), 0.5U/µl Ampligase (Lucigen, A0102K) in 1X Ampligase buffer was added to the samples. The samples were incubated in the gap-filling reaction mix for 5 minutes at 37°C, followed by 40 minutes at 45°C. The samples were then washed twice in PBST and once in hybridization mix (10% Triton X-100 (ThermoFisher, BP151), 2X SSC in water).

#### Rolling circle amplification

The rolling circle amplification (RCA) primer mix (1 µM each of PAB0010, 12, 13, 22, 33, 35, Supplementary Table 2) was then hybridized at room temperature for 15 minutes. The samples were washed twice in hybridization mix for 2 minutes each and once in PBST. The RCA mix containing 0.25 µg/µl non-acetylated BSA (Thermo Fisher Scientific, AM2616), 250 µM dNTP (Thermo Fisher Scientific, R0192), 125 µM 5-(3-Aminoallyl)-dUTP (Thermo Fisher Scientific, AM8439), 5% glycerol, 0.2U/µl EquiPhi29 DNA polymerase (Thermo Fisher Scientific, A39391) in 1X EquiPhi29 buffer was added to the samples and incubated overnight at 30°C in humidified chambers. The following day, samples were washed once in PBST and crosslinked with 50 mM BS(PEG)9 (Thermo Fisher Scientific, 21582) in PBS for 1 hour. Excess BS(PEG)9 was quenched by washing the samples twice with 1 M Tris-HCl pH 8.0 buffer and then incubating at room temperature for 30 minutes. The samples were then washed in PBST.

#### Image acquisition

All tissue imaging was performed using a Nikon Ti2-E inverted microscope, equipped with a PZ-2000 XYZ piezo stage (Applied Scientific Instrumentation) and a Cairn MultiCam beamsplitter and CellCam Kikker 100MT cameras (Cairn Research) using a multibandpass main dichroic and three beamsplitter dichroics to achieve simultaneous collection of four emission channels (see Supplementary Fig. 2a for a list of filters). Fast Z-stack acquisition was achieved using TTL triggering from the exposure out signal of one of the cameras. A 20x 0.8 NA Lambda-D air objective was used for all imaging, with illumination provided by a CoolLED pE-4000 light source. Images were captured with 0.231 µm pixel size in 23-plane Z-stacks with 1 µm spacing and tile overlap of 10%. Microscope hardware was controlled using Micromanager as a GUI interface (59), using Pycromanager (60) to automate the simultaneous acquisition of multiple flowcells.

#### mCherry imaging and photobleaching

The samples were first imaged to detect the fluorescence of mCherry-positive starter neurons using 500 nm excitation light and the same Chroma ZT405/514/635rpc dichroic and Chroma ET590/33m and Chroma ET546/22x emission filters used for sequencing rounds. After imaging, the samples were placed on an aluminum block under a white-light 10,000-lumen LED floodlamp (NATPOW, TS-F42100) overnight to photobleach the mCherry signal to avoid crosstalk with fluorophores during sequencing. A desk fan (Sealey, SFF-04) was used to maintain a consistent sample temperature of under 35°C during photobleaching.

#### Sequencing cell type marker gene transcripts

After mCherry imaging, the samples were equilibrated in hybridization mix and 1 µM LNA endogenous gene sequencing primer (PAB0032, Supplementary Table 2) in hybridization mix was added to the sample and incubated at room temperature for 1 hour. Excess sequencing primer was removed with three washes of hybridization mix at room temperature for two minutes each. The samples were then washed twice in PBST before proceeding to sequencing. Sequencing was performed manually by pipetting Illumina reagents into the flowcells, placing them in a hybridization oven for incubation at 60°C and then aspirating the solution using a vacuum pump. The samples were cooled to room temperature each round before imaging to prevent fluctuations in focus from the infrared-based “Perfect Focus System” of the microscope. The sequencing reagents (HB2, IRM, CRM, and USM) were sourced from HiSeq SBS v4 kits (Illumina, FC-410-1003) or MiSeq Reagent v3 kits (Illumina, MS-102-3003).

For the first cycle of sequencing chemistry, the flowcell was washed twice with 200 µl HB2 buffer (3 min at 60 °C each). Residual thiols were then blocked with 2 × 200 µl iodoacetamide solution incubations (9.3 mg in 2 ml PBST; 3 min at 60 °C). The flowcell was washed once with 200 µl PBST, then washed twice more with 200 µl HB2, all at room temperature. Sequencing was then initiated by introducing 2 × 200 µl IRM (3 min at 60 °C each). Unincorporated dye-conjugated nucleotides were removed with 4 × 200 µl PBST washes (3 min at 60 °C each). Finally, 200 µl USM was applied to reduce photobleaching, and the flowcell was imaged.

Between sequencing cycles, the flowcell was washed with 3 × 200 µl HB2 at room temperature, followed by 2 × 200 µl CRM incubations (3 min at 60 °C each) to cleave the nucleotide terminator groups. The chemistry steps described for the first cycle were then repeated. In total, seven sequencing cycles were completed to read out the entire gene-padlock barcode, using the same wash and incubation parameters at every step.

#### Rabies barcode sequencing

Sequencing of the rabies barcodes took place after the endogenous gene sequencing rounds were complete. The flowcells were washed twice with a stripping buffer of 60% Formamide in 2X SSC buffer at 60°C for 15 minutes to remove the extended gene sequencing primer. The samples were then equilibrated in hybridization mix and 1 µM rabies barcode sequencing primer (PAB0034, Supplementary Table 2) in hybridization mix was added to the samples and incubated at room temperature for 1 hour. Excess sequencing primer was removed with three washes of hybridization mix at room temperature for two minutes each. The samples were then washed twice in PBST before proceeding to sequencing. The same sequencing procedure was carried out as for cell type marker genes, completing 14 rounds of sequencing. As 10 rounds were sufficient to identify unique barcodes (Supplementary Fig. 3) and the error rate of *in situ* sequencing increases over rounds, only the first 10 rounds were included in the analysis.

#### Hybridization rounds for highly expressed genes

The extended barcode sequencing primers were stripped from the samples as described above to prepare them for hybridization probe imaging. The samples were then equilibrated in the hybridization mix, and 1 µM each of a set of hybridization probes with 5’ conjugated fluorophores were hybridized to the samples in two extra rounds. Each round followed the same hybridization and stripping procedure as for the sequencing primers. First, probes designed against the highly-expressed gene padlocks *Slc17a7, Sst, Gad1*, and *Vip* were bound to the samples. These were imaged and then stripped using a solution of 60% formamide in 2X SSC buffer. Then, hybridization probes that targeted a shared sequence on the backbone of all gene padlocks and all rabies barcode padlocks were bound. After adding these ‘anchor’ probes, the tissue was stained with 5 µg/µl DAPI diluted in PBST for 10 minutes, followed by three washes in PBST before proceeding to imaging. First, only the tiles that were imaged for all sequencing rounds were acquired. The acquisition was then repeated, covering the whole brain section to aid with registration to a reference atlas (see below).

#### Sensitivity of barcode detection

To determine the sensitivity of rabies barcode detection, sections of brain tissue infected with the barcoded rabies virus were prepared as detailed above with the following changes: during the reverse transcription step, RT primers for the rabies N, P, M and L genes were added (60 nM per primer, Supplementary Table 3), alongside the pool of endogenous gene RT primers (60 nM per primer) and the rabies barcode padlock (1 µM). Likewise, for the padlock ligation step, padlocks for the rabies N, P, M and L genes were added (60 nM of each padlock, Supplementary Table 3) alongside the endogenous gene padlocks. After RCA was completed, hybridization probes against the rabies gene padlocks (PAB0048, 1 µM) and the rabies barcode padlock (PAB0023, Supplementary Table 2, 1 µM) were added in hybridization mix to the sample and incubated for 1 hour. The sample was washed three times with hybridization mix, then 200 µl USM was added. The samples were then imaged with the same filter set that was used for barcode and high-expressing gene probe imaging.

Rabies-infected cells were identified from the hybridization probe images using the rabies gene signal to segment cells. A difference of Gaussians filter was applied to the images, which were then binarized with a manually selected binarization threshold and binary closing was performed. Candidate rabies-positive cells were then detected on the binarized images. The masks were then filtered based on minimum and maximum area, circularity, elongation, solidity and unbinarized intensity. Each mask was then radially expanded by 5 µm. Rabies barcode spots were detected by finding peaks in the hybridization channel for the barcode probe and the number of barcode spots per rabies-positive cell mask was quantified.

### *In situ* sequencing preprocessing

#### Image projection

Z-stacks were z-projected to calculate the maximum and median intensity projections. The images were flat-field corrected using the average of maximum projections across all rounds and tiles. Since the median projection largely captures diffuse tissue background, all further analyses were done by subtracting the median projection from the maximum projection.

#### Image registration

After projection, images were registered first across successive sequencing rounds, then across channels. A reference tile was manually selected and, for each channel, images from all rounds were median-filtered to reduce noise. Rigid registration parameters between successive rounds of imaging were estimated by performing an iterative grid search over rotation angles using phase correlation to estimate translation. The rotation angle across rounds was only a function of the slide position on the microscope and was constant across tiles. Due to stage inaccuracy, translation parameters could, however, vary from tile to tile and were therefore estimated by phase correlation for each tile independently. To identify aberrant x/y translation shifts, usually at the border of the brain, when a significant part of the image was outside of the sample, shifts across tiles were linearly fitted using a RANSAC regression, and shifts for tiles with residuals above 10 pixels were replaced by the regression value.

To register images across channels and correct chromatic aberration, we then computed the standard deviation projection across registered rounds, generating a reference image for each channel. Images from each channel were tiled into 256 x 256 pixel blocks with 50% overlap. For each block, x/y shifts to the reference channel were calculated by phase correlation. The x and y shifts for each block were then used to estimate the affine transformation between channels. Finally, for each tile, the images for each channel were registered using the affine registration parameters estimated from the reference tile and then further aligned using phase correlation to correct for any channel drift during imaging.

Hybridization acquisitions were registered between channels by calculating the affine transformation between channels on all tiles using a similar method as described above. As these rounds had less signal overlap between channels, registration was performed using blocks of 512 x 512 px with 80% overlap. Each block was binarized using a 0.7 quantile cut-off before registering channels using phase correlation. Finally, affine transform parameters were estimated first between pairs of channels with significant bleedthrough (channels 0 and 1 and channels 2 and 3) and then between the two channels of each group that are the most similar (channels 1 and 3). This procedure was repeated for each tile. Images for each channel were registered using the median affine registration parameters across tiles, and any residual uncorrected translation was estimated by aligning channels to the reference channel using phase correlation.

#### Channel normalization and bleedthrough estimation

Images from sequencing rounds were filtered using a difference of Hanning kernel with radii of 2 and 4 pixels (0.462 and 0.924 µm). To correct for differences in brightness between channels, normalization factors were computed as the 99.99% quantile of pixel intensities for each channel and round, and smoothed by fitting the decrease in intensity across rounds for each channel with an exponential decay. The smoothed normalization factors for round 1 were used to correct brightness across all rounds.

To estimate the expected fluorescence profile of the fluorophores associated with each base, we first detected spots on the standard deviation projection across rounds of the filtered images as local maxima crossing a manually set threshold. We then selected well-isolated spots and extracted fluorescence values of each channel and round in a 2-pixel radius. Scaled K-means clustering (28) was then used to assign each spot to one of four clusters corresponding to the four dye-conjugated nucleotides. The cluster means of the four clusters were computed across rounds, generating a 4 channel x N_rounds_ bleedthrough matrix for each base.

#### Base calling rabies barcodes

Rabies barcodes have unknown sequences and must therefore be basecalled round by round. To do so, putative barcode spots were detected from the registered round/channel images by finding peaks in the mean image across rounds and channels, and the fluorescence trace across rounds and channels was extracted from a disk with a 2-pixel / 0.462 µm radius.

Each spot was assigned to the base with the highest cosine similarity with the bleedthrough matrix for each round. Barcode sequences were constructed from these assignments and the following quality metrics were computed: mean fluorescence intensity across rounds, mean cosine similarity score across rounds, cosine similarity score of the whole sequence, calculated by comparing the observed fluorescence traces (after background subtraction) to synthetic traces generated from the cluster means of the bases which were assigned at each round.

To distinguish genuine barcode spots from bright background autofluorescence spots, spots were first filtered by applying thresholds for the individual quality metrics. To further filter the remaining spots, the quality metrics were fit using a Gaussian Mixture Model (GMM) with two clusters, representing low and high-quality spots. Spots assigned to the high-quality cluster were retained for further analysis, and error correction was performed by merging sequences within a maximum hamming distance of 2.

#### Gene barcode detection using orthogonal matching pursuit

Transcripts of cell-type marker genes were identified by a unique gene barcode sequence encoded in the backbone of gene-specific padlocks. To robustly decode gene spots while accounting for any spatial overlap between them, fluorescence images from gene sequencing rounds were unmixed using a modified version of the orthogonal matching pursuit (OMP) algorithm (28, 61).

First, the expected fluorescence for each gene barcode was computed by concatenating the bleedthrough vectors of the base for each round to generate a gene dictionary. In addition, a set of background vectors representing constant fluorescence across rounds for each channel were appended to the gene dictionary. Fluorescence vectors of individual pixels and elements of the gene dictionary were normalized to unit norm, yielding the normalized fluorescence vector **y** and gene dictionary **G**. OMP approximates the fluorescence **y** as a weighted sum of elements of the dictionary **G** assigned to a given pixel, defined by the support set Λ. Λ is initialized to contain the background vectors and is updated on each iteration to add the gene that best explains the residual fluorescence in the pixel:

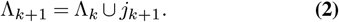

To select gene *j*_*k*+1_ added on a given iteration, we first select the columns of the gene dictionary already included in Λ_*k*_:

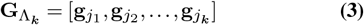

We then fit the fluorescence **y** to estimate the coefficients *β* for each selected gene and background components

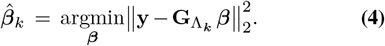

We then compute the residual fluorescence **r**_*k*_

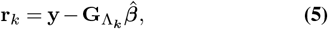

and select the gene *j*_*k*+1_, whose gene vector has the highest absolute dot product with the residual:

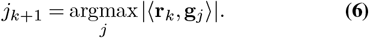

Rounds and channels containing bright signals corresponding to genes included in Λ may not be fully accounted for in the residual fluorescence **r**_*k*_, which could lead to false-positive detection of other genes on subsequent iterations. To control for this, the algorithm above is modified to weigh rounds and channels based on the amount of fluorescence attributed to them (see https://github.com/znamlab/iss-preprocess, based on https://github.com/jduffield65/coppafish).

The iteration continued until a maximum number of components was reached, or the dot product ⟨**r**_*k*_, **g**_*j*_*k*+1⟩ no longer reached a preset threshold. This resulted in a sparse approximation of the fluorescence in every pixel as a combination of a small number of gene-specific components.

The gene coefficients for each gene were then assembled into images and peaks were detected in these gene coefficient images. To distinguish genuine signals from spurious peaks, a spot shape score was assigned to each peak based on the similarity of the signal at that location to a precomputed spot shape image. Any peaks that have a spot score exceeding a manually selected threshold were retained for further analysis.

#### Hybridization spot detection

To detect gene spots, the bleedthrough vectors for each fluorophore were first estimated on a set of reference tiles as described above for sequencing rounds. Peaks were then detected on all tiles, and the mean fluorescence values in each channel at a disk with a 2-pixel / 0.462 µm radius in each spot location were extracted. Fluorescence vectors for each spot were then compared to the bleedthrough matrix and assigned to the probe with the highest cosine similarity.

#### Registration between imaging tiles and cryosections

All image modalities were registered to the first gene round, aligning the gene spots, hybridization spots and DAPI fluorescence images on a tile-by-tile basis by estimating rotation and translation between the target and reference tiles as described above for registration across rounds.

The x/y shifts between adjacent tiles were estimated by phase correlation using the overlap regions. Large outlier shift values or shifts with low correlation maxima were corrected to the median shift along each tile row. These tile shifts were used to combine spot coordinates across tiles into a whole section coordinate system. To avoid duplicate spot detection in overlap regions, spots were only retained if they were closer to the center of the tile on which they were detected than the center of any other tile.

To register the spot coordinates from each section to a common coordinate system and determine their location within the brain, all sections were registered to the Allen CCFv3 10 µm coronal atlas (32). First, stitched images of whole coronal sections were registered to *in situ* sequencing images, which contained only a part of one hemisphere, using masked phase correlation (62). Coronal sections were downsampled, and cutting angles and positioning of slices along the Z-axis were estimated using Deepslice (63). Affine transformation of the slices was performed using Elastix (64) and non-rigid transformation was performed using BigWarp (65), relying on manually selected landmarks. The stitched spot coordinates from each section were then converted to atlas coordinates. The [BrainGlobeAtlas API](https://brainglobe.info/documentation/brainglobe-atlasapi/index.html) (66) was used to define brain area annotations.

The Allen CCDFv3 coordinates were converted to a flattened cortical representation by using the CCF-streamlines package (32). Non-cortical cells situated less than 150 µm from the cortex were projected to their closest neighboring voxel in the cortex. As the thickness of cortical layers varied across the sequenced volume, to plot the cortical depth of cells in flatmap coordinates while respecting layer borders (Fig. 4b and Fig. 5a), layer thickness was rescaled to the average layer thickness across V1.

#### Cell segmentation

Cell nuclei were segmented using Cellpose (67). The “cyto” base model from Cellpose was fine-tuned using manually curated data, with DAPI fluorescence as the nuclei channel and the anchor hybridization probe that binds to all gene spots as a cytoplasmic channel. Cellpose was run on individual planes of DAPI Z-stacks, and the resultant masks were projected to a single plane, removing small masks or masks only found on a single plane and merging any remaining masks based on their spatial overlap.

To assign gene spots to cells, mask images from all tiles in each section were stitched together and dilated by 5 µm to expand the nuclei masks to cover the soma of each neuron. All gene spots within each dilated mask were assigned to it. Assigning all barcode spots present within each dilated mask risks false positives, as rabies barcodes are present in the neuropil as well as in neuronal somata. Instead, barcodes were assigned to cells based on their spatial clustering using a probabilistic assignment model including a Gaussian spatial likelihood function, which considers the distance of a spot to a mask, a background component with a fixed low-density likelihood to account for spots that do not correspond to any specific mask, and a mask spot-count likelihood, which penalizes spot assignments to cells unless justified by the spatial data, encouraging sparseness. Each spot **x**_*i*_ can be assigned to either a mask *j* ∈ {1, …, *M*} or to the background (denoted by − 1). We define a spatial probability density function for a spot to originate from a specific mask or background, defined as

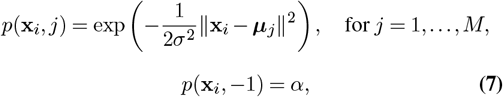

where **x**_*i*_ ∈ ℝ^2^ is the 2D coordinates of spot *i*, ***µ***_*j*_ ∈ ℝ^2^ is the centroid of mask *j, σ* is the spatial scale parameter and *α* is a small constant (e.g. 10^-6^) representing the background likelihood.

We also define a mask spot count probability function

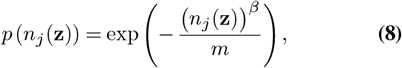

where *n*_*j*_(**z**) is the number of spots assigned to mask *j* under the assignment vector **z**, *m* is the spot count scale factor, and *β* is an exponent that controls how the likelihood penalty scales with spot count. Setting *β <* 1 makes the log-penalty sublinear in *n*_*j*_, so the incremental penalty for each additional spot decreases with every added spot, favoring assigning spots to a small number of masks over splitting them across many masks. Combining the spot-wise assignment and the spot-count terms, the full likelihood for the whole assignment vector **z** is

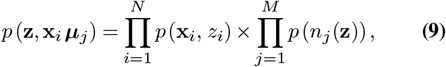

where *N* is the total number of spots and *M* is the total number of cell masks.

Every barcode sequence is handled in isolation, such that the assignment of one barcode does not affect the assignments of other barcodes. Initially, spots falling within a specified maximum distance threshold are assigned to the nearest mask based on Euclidean distance. Spots that do not meet this criterion are assigned to the background. These initial spot assignments are then refined through an iterative optimization process. In each iteration, the algorithm explores potential reassignments of spots to masks by evaluating combinations of spots that are found close together, defined by an inter-spot distance threshold. For each valid combination of spots, the algorithm reassigned spots to a different mask or to the background to maximize the assignment likelihood. The algorithm prioritizes reassignments that yield the highest increase in overall likelihood, thereby iteratively improving the accuracy of spot-to-mask assignments. On each iteration, the algorithm evaluates two scenarios for each spot combination: moving the spots to a new mask and assigning them to the background. The optimization terminates once none of the spots are reassigned on a given iteration or the iteration limit is reached.

#### mCherry cell detection

To detect the nuclei of mCherry-positive starter neurons, mCherry fluorescence was first unmixed using fluorescence from the green (Chroma ET546/22x) channel. A difference of Gaussians filter was applied to the unmixed images, which were then binarized with a manually selected binarization threshold and binary closing was performed. Candidate mCherry nuclei were then detected on the binarized images. The masks were filtered based on minimum and maximum area, circularity, elongation, solidity and unbinarized intensity.

To assign mCherry masks to rabies-infected cells, the reference-aligned tile coordinates were used to make full-section stitched images of the mCherry masks and rabies spots coordinates. Each mCherry mask was first checked for any rabies spots that overlapped with it. If barcode spots were found in the mCherry mask, the cell to which the most abundant barcode belonged was checked to find all of its barcode spots. If the percentage of spots from that cell found within the mCherry mask was above a threshold (conservatively 10% of the total spots in the rabies-infected cell), and the centroid of the spots overlapped with the mCherry mask, then the mCherry mask was assigned to that cell.

#### Cell type classification

Single cell gene expression data was analyzed using the scanpy single-cell analysis toolkit (68). First, cell-type clusters were defined for high-quality cells in a reference dataset collected from four coronal sections covering the visual cortex in a separate brain. High-quality cortical cells were selected as cells with >=4 unique genes per cell and >=30 gene counts per cell. Transcript counts in each cell were normalized and log1p-transformed (natural log with a pseudo-count of 1). The first 30 principal components were then used to compute the nearest-neighbors distance matrix and a neighborhood graph of the cells with the local neighborhood size set to 10. Cells were then clustered into subgroups using the Leiden algorithm (69). The resulting clusters were manually annotated as cell types by referencing the expression of marker genes in the gene panel to a V1 SMARTseq scRNAseq dataset (12) and by comparing the spatial location of each cell cluster with known cortical layers. Mean gene expression values were calculated per cluster.

For each cell *j* in the rabies *in situ* sequencing dataset, we calculated the Pearson correlation coefficient *r*_*jk*_ = corr(**x**_*j*_, ***µ***_*k*_) between the cell’s log-normalized expression vector **x**_*j*_ and the log-mean expression vector ***µ***_*k*_ of reference cluster *k*. For the analyses in Fig. 5a-g, excitatory rabies-infected cells were selected based on the identity of their highest correlation cluster. For the analyses in Fig. 5h-j, rabies-infected cells were classified as inhibitory if the cluster yielding their maximum correlation corresponded to one of the inhibitory reference clusters *Pvalb, Sst, Vip*, or *Lamp5* and if they satisfied all three of the following thresholds: a Pearson correlation coefficient of 0.3, a minimum gene count of 3, and a minimum nearest-neighbor cluster label agreement of 0.3.

Clusters were visualized using Uniform Manifold Approximation and Projection (UMAP) (70) with a minimum distance between embedded points of 0.05. Scanpy’s ingest function was used to project all *in situ* sequenced cells onto the PCA space of the high-quality cell reference data, followed by k-nearest-neighbor classification and UMAP embedding transfer for visualization of reference clusters (Fig. 2h, i).

### Data analysis

#### Estimation of the proportion of cells present in the sequenced volume

To determine the proportion of starter cells contained in the *in situ* sequencing volume (Supplementary Fig. 4a), sections adjacent to the sequenced dataset were imaged and analyzed by following the mCherry imaging and mCherry cell detection steps described above. To estimate the number of presynaptic cells included in the dataset (Supplementary Fig. 4b), the *in situ* sequenced volume was registered to the Allen CCFv3 reference atlas as described above. The sequenced volume was then superposed onto bulk rabies tracing data (Fig. 1h-j) by aligning the manually determined injection centers of the two experiments and the proportion of rabies-infected cells within this volume in bulk tracing data was quantified. For illustration, we plotted the medio-lateral and dorso-ventral position of cells within 400 µm of the central coronal plane over the atlas borders of this coronal plane (Supplementary Fig. 4b top) and the antero-posterior and medio-lateral position of all cells overlaid on a dorsal view of the atlas borders (Supplementary Fig. 4b bottom). The sequencing sections were not perfectly coronal and thalamic cells present in the volume (orange) appear anterior to the cortical cells.

#### Probability of barcode transmission between starter cells

The probability that a given starter neuron transmits its barcode to another starter cell (Fig. 1g), *p*, was estimated as a function of the number of neurons infected by each starter cell, *n*, and the proportion of neurons infected by the starter AAVs, *α*:

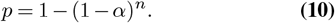

To convert between the proportion of starter neurons and cell density, the overall density of neurons in the visual cortex was estimated as 150,000 cells per mm^3^ (71, 72).

#### Library barcode abundance

To compare the sequences of barcodes found *in situ* to those found in the viral library (Fig. 3b, Supplementary Fig. 6a), we first determined the abundance distribution of barcodes in the library by grouping barcodes by their number of unique reads (sequences with the same barcode but different UMIs), and plotting the total proportion of the library represented in each of these abundance bins (Fig. 3b, black). Given that only a few barcodes have high abundance (Fig. 1c), they collectively represented only a small fraction of total reads in the library, and most reads of the library consisted of barcodes representing each about 10^-5^ of the library. We then matched the 2,158 barcodes found *in situ* and the same number of randomly generated barcodes to the library barcodes and plotted the distribution of the abundance in the library of those matches (Fig. 3b, blue lines).

To determine the excess number of high abundance barcodes among those present in multiple starter cells (Supplementary Fig. 6b), the kernel density estimate of the distribution of barcode proportion of unique reads in the library (black, Supplementary Fig. 6a) was subtracted from that of barcodes present in multiple starters (pink, Supplementary Fig. 6a). Then, the area under the curve of this difference (pink, Supplementary Fig. 6b) was numerically integrated between 1.49 x 10^-4^ (where the difference exceeded 0) and 10^-2^ to obtain the proportion of excess barcodes, and multiplied by the total number of barcodes found in multiple cells (n=122), to give the final estimate of 8/122 barcodes.

#### Starter cell mCherry fluorescence analysis

The H2B-mCherry intensity (Fig. 3f) was measured as the average fluorescence intensity of the red channel inside each starter cell mask. The logarithm of the intensity was then correlated to the logarithm of the number of presynaptic cells detected for each starter + 1.

#### Relative spatial location of presynaptic neurons

Position of the presynaptic cells relative to their starter cells (Fig. 4f top) were calculated using flatmap coordinates. The distribution of relative medio-lateral location (Fig. 4g) was the kernel density estimate of the difference in medio-lateral position only with a bandwidth of 50 µm. To estimate a null distribution assuming random connectivity, we shuffled the barcodes of presynaptic cells but not those of starter cells. This procedure preserves the number of presynaptic cells per starter and the spatial distribution of starter and presynaptic cells. A single example shuffled sample was used for the scatter plot of Fig. 4f (bottom). 10,000 bootstrap samples were used to calculate the confidence interval of medio-lateral distance distribution (Fig. 4g dashed line). The median distance to the starter cells was computed in flatmap coordinates using the 3 dimensions and compared to that of the shuffled samples (see Results).

#### Layer- and cell-type-specific connectivity

The analysis of local excitatory connectivity in Fig. 5a-g included all putative local excitatory connections by removing cells classified as inhibitory and connections from presynaptic cells that were more than 1 mm away from their starter cell. Only starter cells with at least one local presynaptic cell were included for analysis. Input fractions (Fig. 5b-e) were calculated as the number of connections from each layer divided by the total number of local excitatory connections for each starter and averaged across starters in each layer. Output fractions (Fig. 5f,g) were computed by dividing the number of connections (Fig. 5b, left) by the sum of the number of connections per presynaptic layer.

To identify over- and under-represented connections, we shuffled the barcodes across presynaptic neurons, keeping their layer identity and spatial location unchanged, repeating the shuffling procedure 10,000 times. Effect sizes were computed as the ratio of the observed input and output fractions and the means of the shuffled samples (Fig. 5e,g, circle diameter and colormap sign). To estimate p-values, we computed the log effect size for individual shuffle samples and determined the quantile *q* of the effect size distribution at 0. We then calculated the uncorrected two-sided p-value as 2 min {*q*, 1− *q*}. These p-values from the bootstrap distribution were then corrected for a false discovery rate of 5% using Benjamini–Hochberg correction (Fig. 5e, colormap absolute value).

In the analysis of connectivity between inhibitory cell types (Fig. 5i), input fractions were calculated as the number of connections from each cell type divided by the total number of inhibitory connections for each starter and averaged across starters.

Confidence intervals for input fractions (Fig. 5c,d,j) were computed by bootstrap resampling starter cells with replacement 10,000 times as the [2.5-97.5] percentiles of the input fraction distributions.

#### Retinotopic location estimation

To relate long-range connectivity patterns to retinotopic organization of the visual cortex (Fig. 6), azimuth maps from Waters et al. (44) were averaged and aligned to the top view projection. Each starter cell was assigned an azimuth location by projecting its coordinates onto the top view before querying the averaged map. To display the azimuth map in flatmap coordinates, we found the surface 3D voxel of each pixel on the top view projection (using the ‘view_lookup’ of the ccf_streamlines package) and mapped it to the closest flatmap coordinates. Missing pixels were then filled by bicubic interpolation.

The average map of starter ML position (Fig. 6c) was computed as the 2D Gaussian weighted average (*σ*=200 µm) of data in Fig. 6a. This map was clipped using the concave hull of the presynaptic cells’ position to avoid extrapolation outside of the data range. A transparency mask proportional to the total weight of the 2D Gaussian was applied to indicate the number of data points at each map location.

Figure 1 Schematic of the EC-GMR and measurement setup. The EC-GMR consists of a 555nm period GMR 150 nm deep Si^3^N^4^ grating on glass coating in a layer of ITO. For measuring the resonance wavelength a broadband white light source is collimated and the polarisation setup by a Glan Thompson polariser mounted in a rotation mount. The incident light is focused into the back focal plane of the 10x objective by the field lens producing a collimated incident beam at the EC-GMR. The reflected light from the EC-GMR is fibre coupled into the spectrometer. For electrochemical measurement the ITO surface is used as the working electrode, with the Ag/AgCl reference electrode and Pt counter electrode placed in solution above the EC-GMR.

Figure 2 Simulation of EC-GMR resonance wavelength with ITO thickness a Simulated TE reflectance spectra of EC-GMR with different thicknesses of ITO from 0 to 100 nm. b Electric field distribution of the TE mode of the EC-GMR coated with 60nm of ITO. c Simulated TE reflectance spectra of EC-GMR for increasing refractive indexes of a 1nm accumulation region at the ITO media interface. d Peak resonance wavelength against accumulation refractive index change, R^2^= 0.98.

Figure 3 EC-GMR Grating Structure and Resonance Modulation by Bias Voltage: a SEM image of EC-GMR grating which has been cleaved through to expose the profile of the grating. ITO thickness is 62 nm. inset SEM image of EC-GMR grating grooves. b Reflectance spectra of the TE mode for the Si3N4 grating, 20 nm ITO EC-GMR grating and 60 nm EC-GMR grating. Reflectance has been normalised to an Ag mirror. c Hyperspectral image of resonance wavelengths for Si3N4 GMR grating (left) and EC-GMR with 60 nm thick ITO layer (right). The resonance wavelength has been normalised between 0 and 1 for both images. Image area is 181×181 µm. d Change in resonance wavelength Δλ^res^ for 60 nm EC-GMR in 100 mM KPi buffer with increasing bias voltage from 0 to −1. Reflectance spectra shown in inset.

---

**Supplementary Figure 1.**
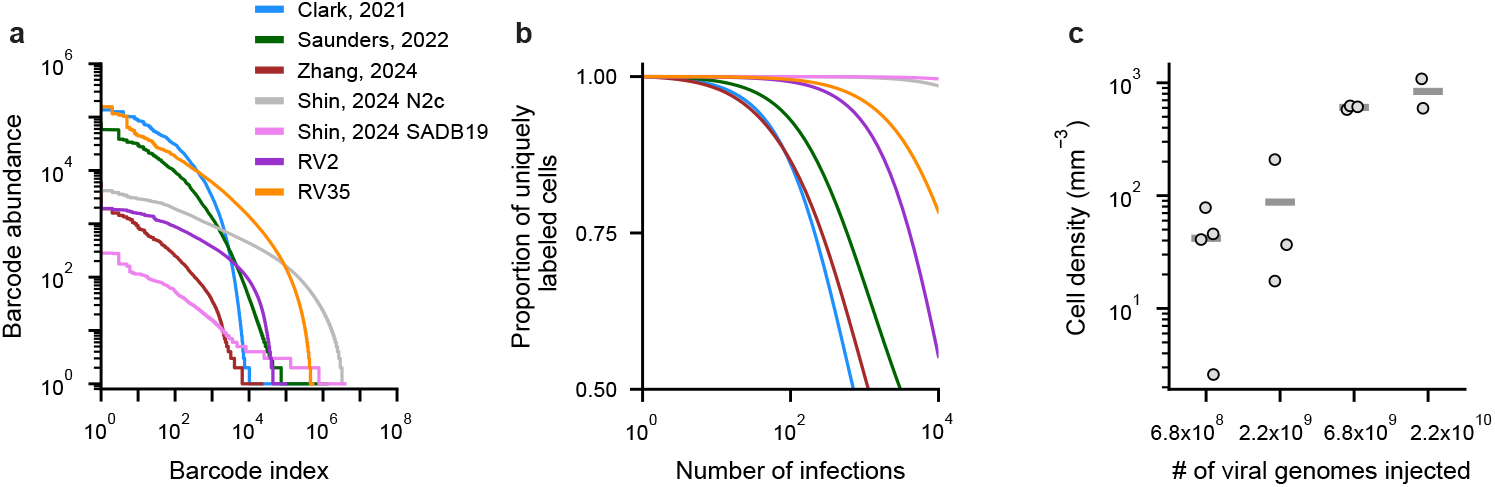
Optimizing barcode library diversity and starter cell density. **a**, Barcode abundance distribution of viral libraries as in Fig. 1c, for previously described barcoded rabies libraries, a lower diversity barcoded rabies library produced before optimizing viral packaging (RV2), and a high diversity barcoded rabies library used for *in situ* sequencing experiments (RV35). **b**, Proportion of uniquely labeled starter neurons as a function of the number of independent infections as in Fig. 1d. Libraries can label 95% of cells uniquely for the following number of independent infection events: Clark 2021 – 33 cells, Saunders 2022 – 69 cells, Zhang 2024 – 28 cells, Shin 2024 N2c – 35,728 cells, Shin 2024 SADB19 – 151,931 cells, RV2 – 630 cells, RV35 – 1,320 cells. The barcode design of Shin et al., which contains common sequences across barcode regions, is susceptible to template switching (45, 73), potentially inflating estimates of library diversity. **c**, Density of tdTomato-positive cells in the visual cortex of Ai14(RCL-tdT)-D Cre reporter mice after tail vein injections of serial dilutions of AAV-PHP.eB-Cre. Bars represent mean density (N = 4 mice for 6.8 x 10^8^ vg/ml, N = 3 mice for 2.2 x 10^9^, 6.8 x 10^9^ vg/ml, N = 2 for 2.2 x 10^10^ vg/ml).

**Supplementary Figure 2.**
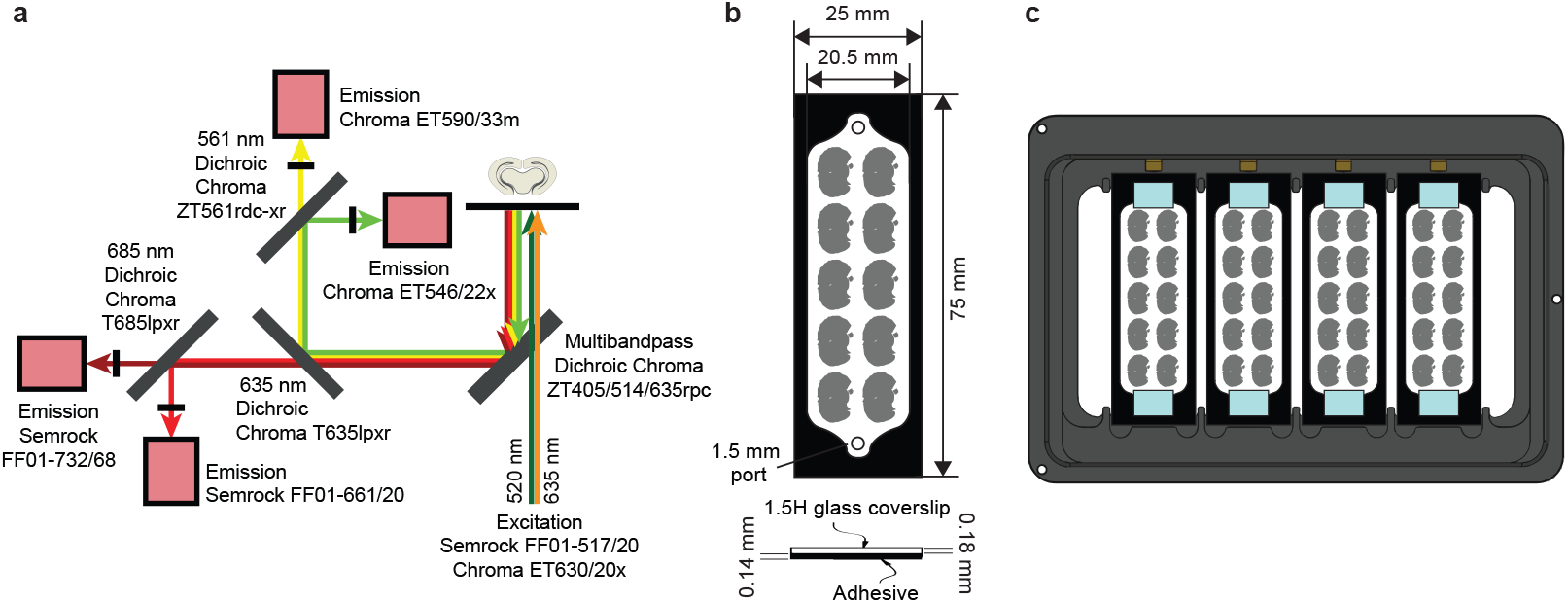
Setup for high-throughput *in situ* sequencing. **a**, Diagram of the light path of the *in situ* sequencing microscope for imaging four channels simultaneously. **b**, Diagram of a microscope slide sized flowcell with a top-down view above and a side-on view below showing mouse coronal brain sections to scale. **c**, Diagram of the flowcells in the 4-slide microscope stage attachment used for the sequencing microscopes.

**Supplementary Figure 3.**
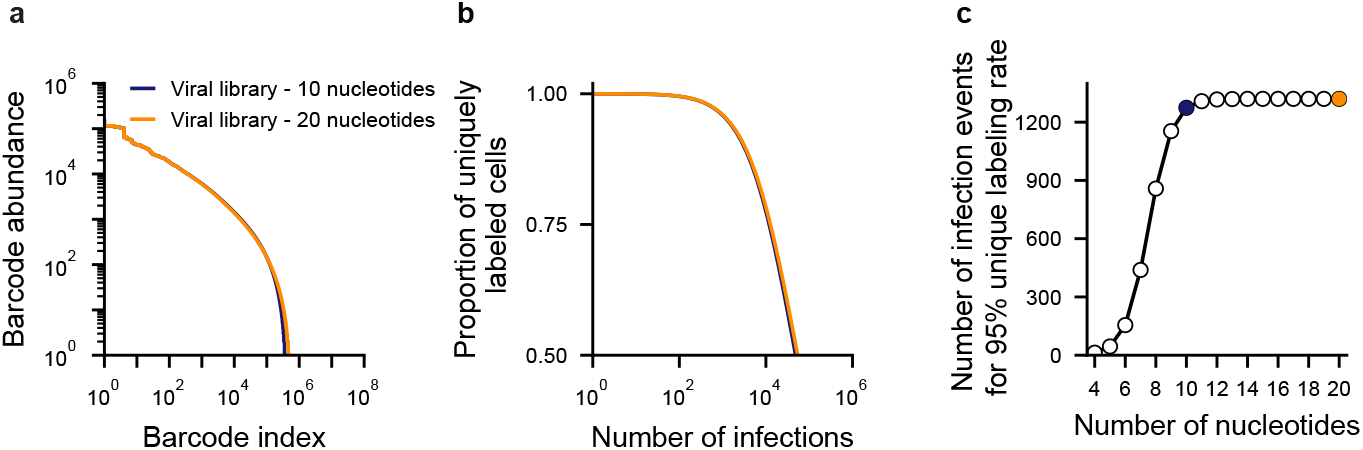
Sequencing 10 nucleotides of the viral barcode is sufficient to uniquely identify infected cells. **a**, Barcode abundance distribution of plasmid and viral libraries as in Fig. 1c, quantified using the full barcode sequence (orange) and the first 10 nucleotides of the barcode (blue). **b**, Proportion of uniquely labeled starter neurons as a function of the number of independent infections, as in Fig. 1d. **c**, Maximum number of cells that can be labeled with at least 95% of uniquely labeled cells as a function of barcode length. Colored dots indicate the full library length (20 nucleotides, orange) and the length sequenced *in situ* (10 nucleotides, blue)

**Supplementary Figure 4.**
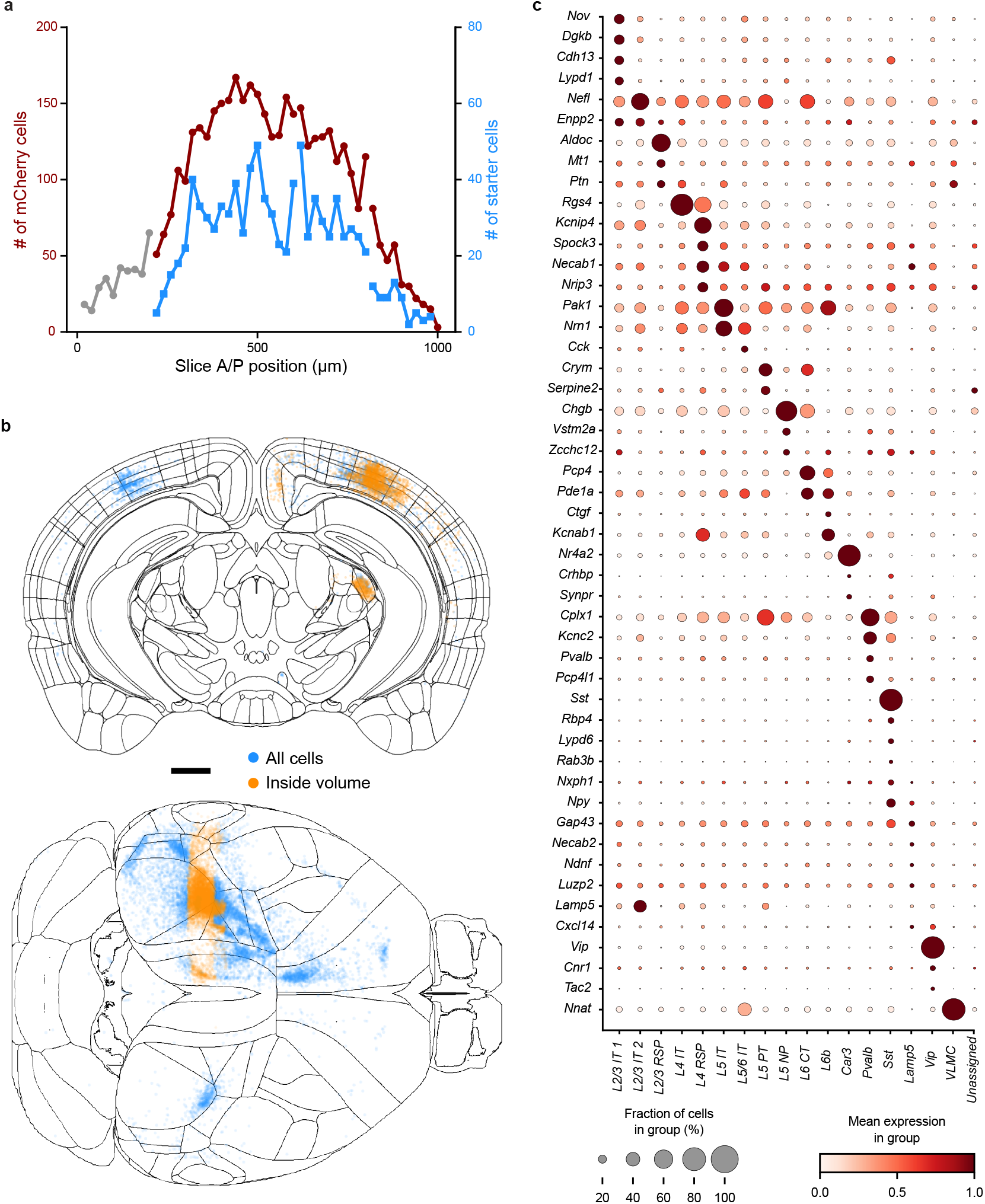
Proportion of cells captured within the sequenced volume and gene expression across clusters. **a**, Number of mCherry cells (circles) detected in each section in sequenced sections (red) or non-sequenced sections (grey) as a function of the antero-posterior section position. The number of rabies-infected starter cells (blue square, right y-axis) found in each section follows the distribution of the total number of mCherry cells per section (infected and uninfected). Line breaks indicate tissue sections in different flowcells. **b**, Coronal (top) and dorsal (bottom) projections of bulk-labeled rabies cell locations. The extent of the *in situ* sequencing volume was centered on the bulk-labeled injection site, and cells were classified as being within the extent of the volume (orange, 8,157 cells) or outside of the volume (blue, 10,160 cells). Scale bar 1 mm **c**, Dotplot of mean gene expression across all cell type clusters for a selection of 49 marker genes from the total panel of 81 genes with cluster-specific expression. Mean normalized and log1p-transformed expression of each gene was rescaled to 1 across cell-type clusters.

**Supplementary Figure 5.**
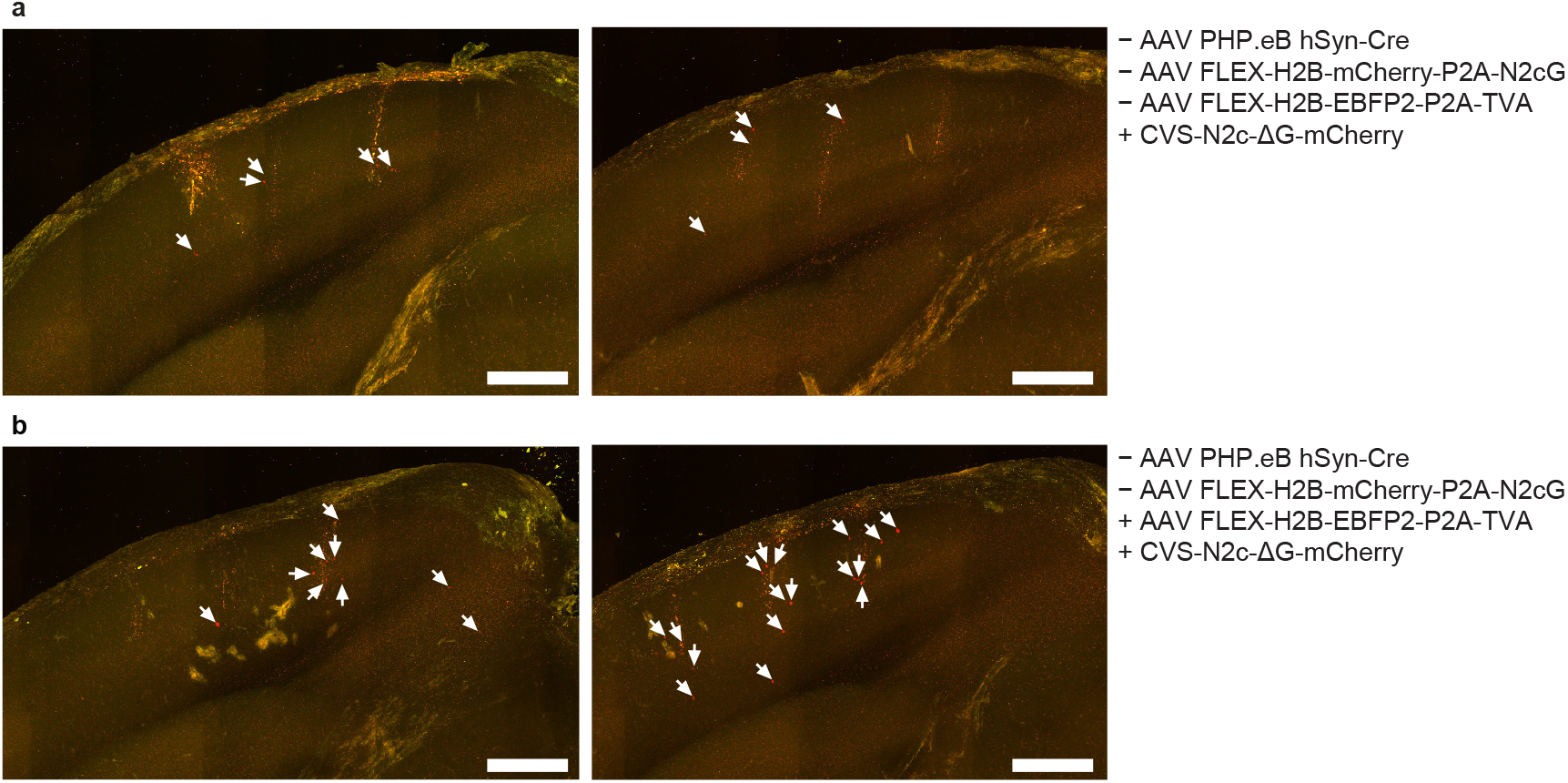
Control experiments for direct infection by rabies viruses from glycoprotein-coated rabies contamination or AAV-TVA leak. **a**, Serial two-photon maximum coronal z-projections displaying 1 mm thickness of tissue centered on the injection sites of barcoded rabies in two brains. No TVA/glycoprotein/Cre virus injections were performed. **b**, Maximum z-projection through 1 mm of tissue from injection sites of both barcoded rabies and AAV-TVA in two brains. White arrows show the location of all detected rabies-infected cells. Scale bar – 500 µm.

**Supplementary Figure 6.**
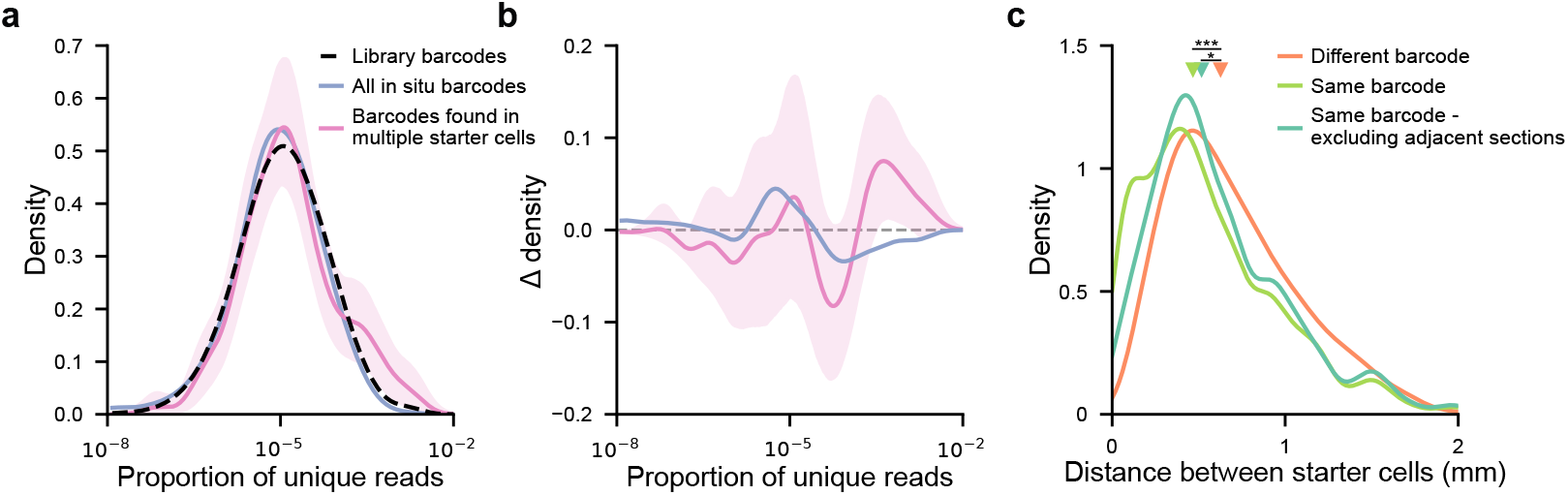
Barcodes found in multiple starters are more abundant in the library and cluster spatially *in situ*. **a**, Kernel density estimates of the representation of barcodes as a proportion of unique reads in the viral library for all library barcodes (black, data as in Fig. 3b), all barcodes found in starter cells *in situ* (blue) or barcodes found in multiple starter cells *in situ* (pink). Shading indicates 95% confidence intervals computed by bootstrap resampling of barcodes found in multiple starter cells. **b**, Difference between the kernel density estimates for all barcodes or barcodes found in multiple starter cells and that of the library barcodes. Shading indicates 95% confidence intervals, computed by bootstrap resampling of barcodes found in multiple starter cells. **c**, The distance between starter cells containing the same barcode (light green, median 0.47 mm, 122 barcodes, 277 starter cells, 297 distances) was smaller than between these starter cells and starter cells with a different barcode (orange, median = 0.63 mm, 564 barcodes, 579 starter cells, 173,384 distances, p<0.001). This difference remained significant after excluding starter cells with the same barcodes found in adjacent sections (turquoise, median = 0.52 mm, 94 barcodes, 135 starter cells, 239 distances, p=0.028) and is therefore not due to duplicate detection of starter cells. Shading indicates 95% confidence interval, triangles indicate medians.

**Supplementary Table S1.**
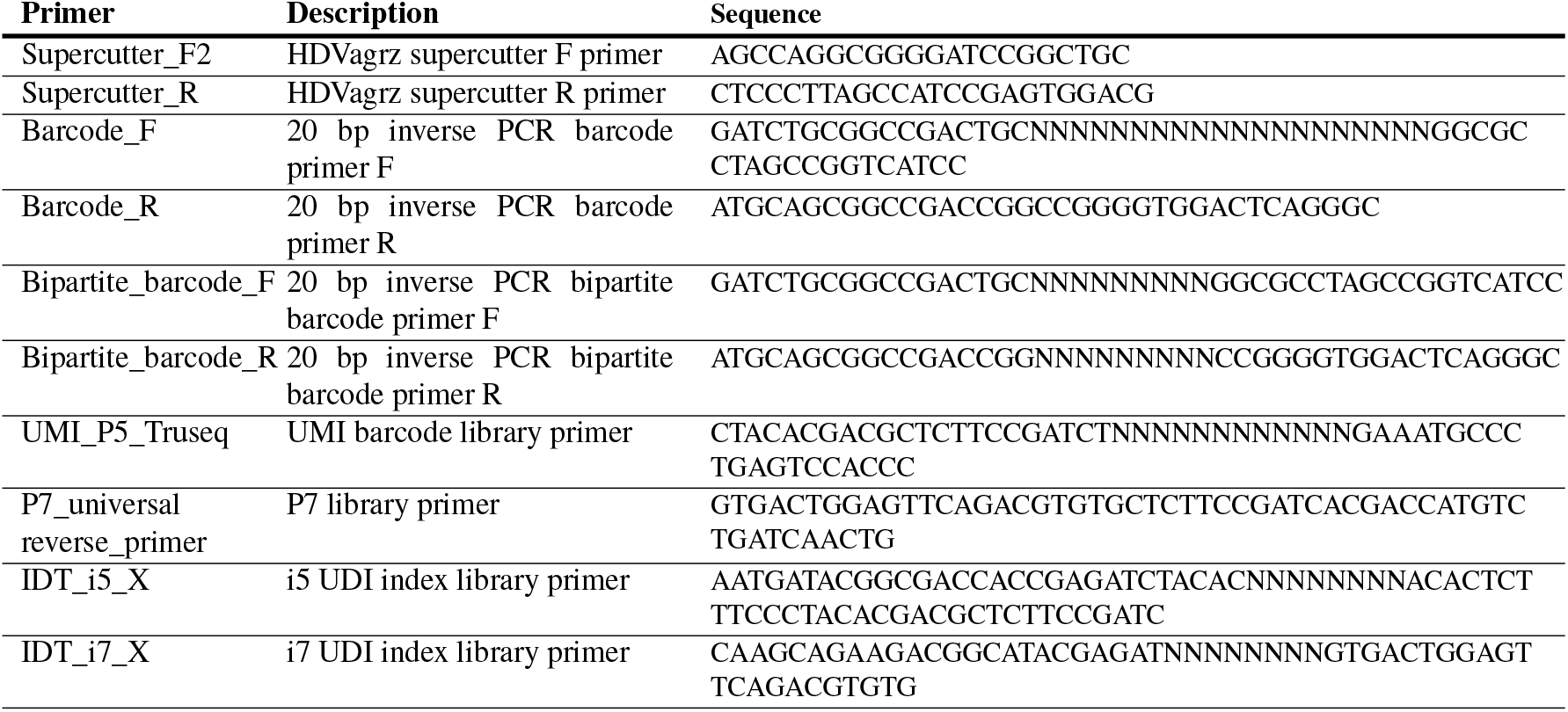
Library cloning and sequencing primers.

**Supplementary Table S2.**
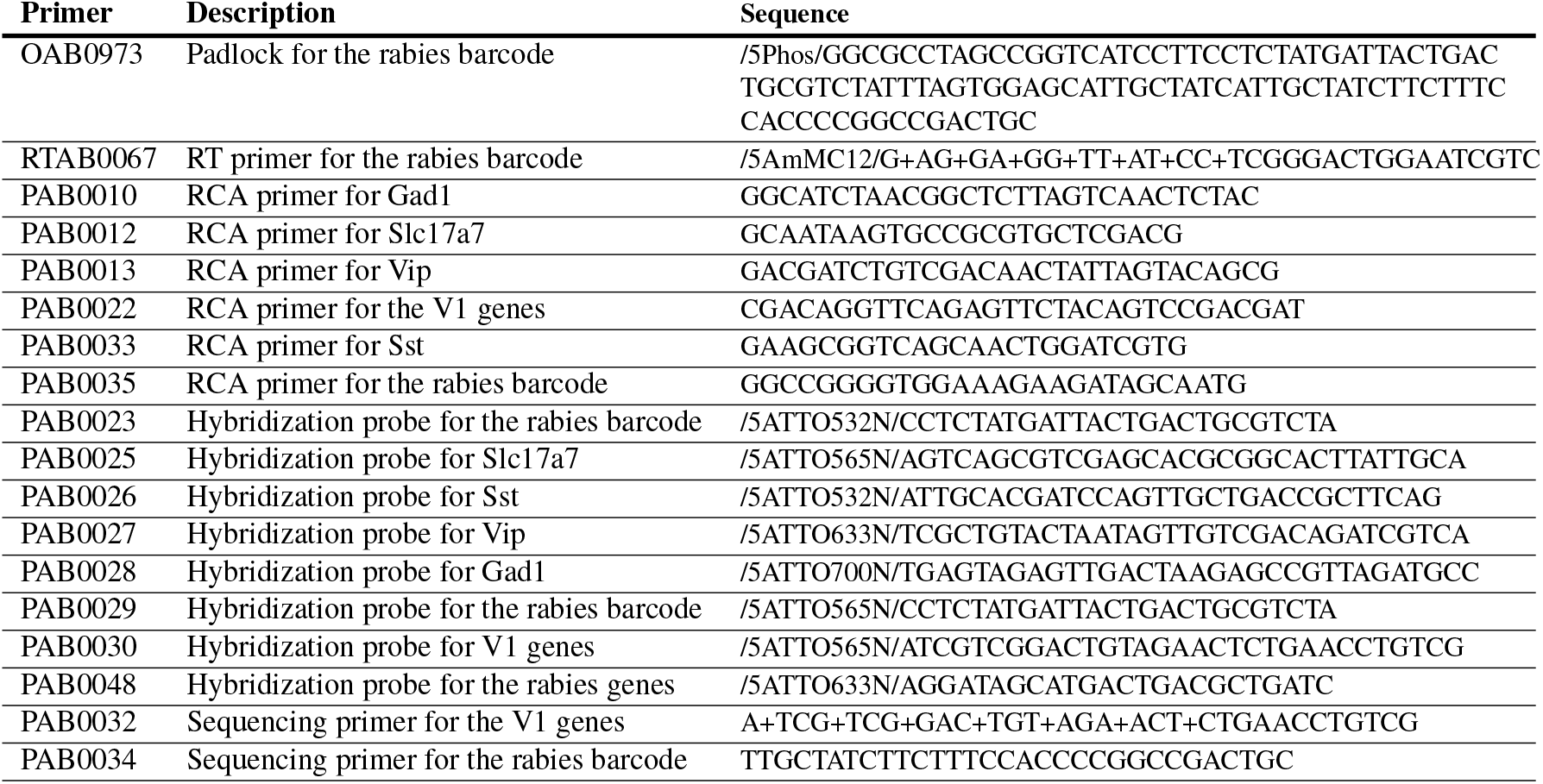
*In situ* sequencing oligonucleotides.

## Notes

### Competing Interest Statement

The authors have declared no competing interest.

## References

1. Zador, A.M., Dubnau, J., Oyibo, H.K., Zhan, H., Cao, G., and Peikon, I.D. (2012). Sequencing the connectome. PLOS Biology, 10(10):1–7. doi: 10.1371/journal.pbio.1001411.

2. Wickersham, I.R., Finke, S., Conzelmann, K.K., and Callaway, E.M. (2007). Retrograde neuronal tracing with a deletion-mutant rabies virus. Nature Methods, 4(1):47–49. doi: 10.1038/nmeth999.

3. Wall, N.R., Wickersham, I.R., Cetin, A., De La Parra, M., and Callaway, E.M. (2010). Monosynaptic circuit tracing in vivo through Cre-dependent targeting and complementation of modified rabies virus. Proceedings of the National Academy of Sciences, 107(50):21848–21853. doi: 10.1073/pnas.1011756107.

4. Saunders, A., Huang, K.W., Vondrak, C., Hughes, C., Smolyar, K., Sen, H., Philson, A.C., Nemesh, J., Wysoker, A., Kashin, S., et al. (2022). Ascertaining cells’ synaptic connections and RNA expression simultaneously with barcoded rabies virus libraries. Nature Communications, 13(1):6993. doi: 10.1038/s41467-022-34334-1.

5. Zhang, A., Jin, L., Yao, S., Matsuyama, M., van Velthoven, C.T., Sullivan, H.A., Sun, N., Kellis, M., Tasic, B., Wickersham, I., et al. (2024). Rabies virus-based barcoded neuroanatomy resolved by single-cell RNA and in situ sequencing. eLife, 12:RP87866. doi: 10.7554/eLife.87866.

6. Lin, Z., Ophir, O., Trimbuch, T., Brokowski, B., Tibi, M., Pavletsov, P., Schmidt, H., Rosenmund, C., Hochgerner, H., and Zeisel, A. (2024). Brain-wide monosynaptic connectivity mapping with ROInet-seq. bioRxiv preprint. doi: 10.1101/2024.04.04.588058.

7. Shin, D., Urbanek, M.E., Larson, H.H., Moussa, A.J., Lee, K.Y., Baker, D.L., Standen-Bloom, E., Ramachandran, S., Bogdanoff, D., Cadwell, C.R., et al. (2024). High-complexity barcoded rabies virus for scalable circuit mapping using single-cell and single-nucleus sequencing. bioRxiv preprint. doi: 10.1101/2024.10.01.616167.

8. Bae, J.A., Baptiste, M., Baptiste, M.R., Bishop, C.A., Bodor, A.L., Brittain, D., Brooks, V., Buchanan, J., Bumbarger, D.J., Castro, M.A., et al. (2025). Functional connectomics spanning multiple areas of mouse visual cortex. Nature, 640(8058):435–447. doi: 10.1038/s41586-025-08790-w.

9. Shapson-Coe, A., Januszewski, M., Berger, D.R., Pope, A., Wu, Y., Blakely, T., Schalek, R.L., Li, P.H., Wang, S., Maitin-Shepard, J., et al. (2024). A petavoxel fragment of human cerebral cortex reconstructed at nanoscale resolution. Science, 384(6696):eadk4858. doi: 10.1126/science.adk4858.

10. Dorkenwald, S., Matsliah, A., Sterling, A.R., Schlegel, P., Yu, S.c., McKellar, C.E., Lin, A., Costa, M., Eichler, K., Yin, Y., et al. (2024). Neuronal wiring diagram of an adult brain. Nature, 634(8032):124–138. doi: 10.1038/s41586-024-07558-y.

11. Tasic, B., Menon, V., Nguyen, T.N., Kim, T.K., Jarsky, T., Yao, Z., Levi, B., Gray, L.T., Sorensen, S.A., Dolbeare, T., et al. (2016). Adult mouse cortical cell taxonomy by single cell transcriptomics. Nature Neuroscience, 19(2):335–346. doi: 10.1038/nn.4216.

12. Tasic, B., Yao, Z., Graybuck, L.T., Smith, K.A., Nguyen, T.N., Bertagnolli, D., Goldy, J., Garren, E., Economo, M.N., Viswanathan, S., et al. (2018). Shared and distinct transcriptomic cell types across neocortical areas. Nature, 563(7729):72–78. doi: 10.1038/s41586-018-0654-5.

13. Zeisel, A., Hochgerner, H., Lönnerberg, P., Johnsson, A., Memic, F., van der Zwan, J., Häring, M., Braun, E., Borm, L.E., La Manno, G., et al. (2018). Molecular architecture of the mouse nervous system. Cell, 174(4):999–1014.e22. doi: 10.1016/j.cell.2018.06.021.

14. Zhang, M., Eichhorn, S.W., Zingg, B., Yao, Z., Cotter, K., Zeng, H., Dong, H., and Zhuang, X. (2021). Spatially resolved cell atlas of the mouse primary motor cortex by MERFISH. Nature, 598(7879):137–143. doi: 10.1038/s41586-021-03705-x.

15. Yao, S., Wang, Q., Hirokawa, K.E., Ouellette, B., Ahmed, R., Bomben, J., Brouner, K., Casal, L., Caldejon, S., Cho, A., et al. (2023). A whole-brain monosynaptic input connectome to neuron classes in mouse visual cortex. Nature Neuroscience, 26(2):350–364. doi: 10.1038/s41593-022-01219-x.

16. Economo, M.N., Viswanathan, S., Tasic, B., Bas, E., Winnubst, J., Menon, V., Graybuck, L.T., Nguyen, T.N., Smith, K.A., Yao, Z., et al. (2018). Distinct descending motor cortex pathways and their roles in movement. Nature, 563(7729):79–84. doi: 10.1038/s41586-018-0642-9.

17. Scala, F., Kobak, D., Bernabucci, M., Bernaerts, Y., Cadwell, C.R., Castro, J.R., Hartmanis, L., Jiang, X., Laturnus, S., Miranda, E., et al. (2021). Phenotypic variation of transcriptomic cell types in mouse motor cortex. Nature, 598(7879):144–150. doi: 10.1038/s41586-020-2907-3.

18. Peikon, I.D., Kebschull, J.M., Vagin, V.V., Ravens, D.I., Sun, Y.C., Brouzes, E., Corrêa, I.R., Bressan, D., and Zador, A.M. (2017). Using high-throughput barcode sequencing to efficiently map connectomes. Nucleic Acids Research, 45(12):e115–e115. doi: 10.1093/nar/gkx292.

19. Oyibo, H., Cao, C., Ferrante, D.D., Zhan, H., Koulakov, A., Enquist, L., Dubnau, J., and Zador, A. (2018). A computational framework for converting high-throughput DNA sequencing data into neural circuit connectivity. bioRxiv preprint. doi: 10.1101/244079.

20. Reardon, T.R., Murray, A.J., Turi, G.F., Wirblich, C., Croce, K.R., Schnell, M.J., Jessell, T.M., and Losonczy, A. (2016). Rabies virus CVS-N2cΔG strain enhances retrograde synaptic transfer and neuronal viability. Neuron, 89(4):711–724. doi: 10.1016/j.neuron.2016.01.004.

21. Schnell, M., Mebatsion, T., and Conzelmann, K. (1994). Infectious rabies viruses from cloned cDNA. The EMBO Journal, 13(18):4195–4203. doi: 10.1002/j.1460-2075.1994.tb06739.x.

22. Conzelmann, K.K. and Schnell, M. (1994). Rescue of synthetic genomic RNA analogs of rabies virus by plasmid-encoded proteins. Journal of Virology, 68(2):713–719. doi: 10.1128/jvi.68.2.713-719.1994.

23. Ghanem, A., Kern, A., and Conzelmann, K.K. (2012). Significantly improved rescue of rabies virus from cDNA plasmids. European Journal of Cell Biology, 91(1):10–16. doi: 10.1016/j.ejcb.2011.01.008.

24. Kebschull, J.M. and Zador, A.M. (2018). Cellular barcoding: lineage tracing, screening and beyond. Nature Methods, 15(11):871–879. doi: 10.1038/s41592-018-0185-x.

25. Clark, I.C., Gutiérrez-Vázquez, C., Wheeler, M.A., Li, Z., Rothhammer, V., Linnerbauer, M., Sanmarco, L.M., Guo, L., Blain, M., Zandee, S.E.J., et al. (2021). Barcoded viral tracing of single-cell interactions in central nervous system inflammation. Science, 372(6540). doi: 10.1126/science.abf1230.

26. Chatterjee, D., Marmion, D.J., McBride, J.L., Manfredsson, F.P., Butler, D., Messer, A., and Kordower, J.H. (2022). Enhanced CNS transduction from AAV.PHP.eB infusion into the cisterna magna of older adult rats compared to AAV9. Gene Therapy, 29(6):390–397. doi: 10.1038/s41434-021-00244-y.

27. Sun, Y.C., Chen, X., Fischer, S., Lu, S., Zhan, H., Gillis, J., and Zador, A.M. (2021). Integrating barcoded neuroanatomy with spatial transcriptional profiling enables identification of gene correlates of projections. Nature Neuroscience, 24(6):873–885. doi: 10.1038/s41593-021-00842-4.

28. Bugeon, S., Duffield, J., Dipoppa, M., Ritoux, A., Prankerd, I., Nicoloutsopoulos, D., Orme, D., Shinn, M., Peng, H., Forrest, H., et al. (2022). A transcriptomic axis predicts state modulation of cortical interneurons. Nature, 607(7918):330–338. doi: 10.1038/s41586-022-04915-7.

29. Liu, Z., Chen, O., Wall, J.B.J., Zheng, M., Zhou, Y., Wang, L., Ruth Vaseghi, H., Qian, L., and Liu, J. (2017). Systematic comparison of 2A peptides for cloning multi-genes in a polycistronic vector. Scientific Reports, 7(1):2193. doi: 10.1038/s41598-017-02460-2.

30. Kim, E.J., Jacobs, M.W., Ito-Cole, T., and Callaway, E.M. (2016). Improved monosynaptic neural circuit tracing using engineered rabies virus glycoproteins. Cell Reports, 15(4):692–699. doi: 10.1016/j.celrep.2016.03.067.

31. Jia, F., Li, L., Liu, H., Lv, P., Shi, X., Wu, Y., Ling, C., and Xu, F. (2021). Development of a rabies virus-based retrograde tracer with high trans-monosynaptic efficiency by reshuffling glycoprotein. Molecular Brain, 14(1):109. doi: 10.1186/s13041-021-00821-7.

32. Wang, Q., Ding, S.L., Li, Y., Royall, J., Feng, D., Lesnar, P., Graddis, N., Naeemi, M., Facer, B., Ho, A., et al. (2020). The Allen mouse brain Common Coordinate Framework: a 3D reference atlas. Cell, 181(4):936–953.e20. doi: 10.1016/j.cell.2020.04.007.

33. Brown, D., Altermatt, M., Dobreva, T., Chen, S., Wang, A., Thomson, M., and Gradinaru, V. (2021). Deep parallel characterization of AAV tropism and AAV-mediated transcriptional changes via single-cell RNA sequencing. Frontiers in Immunology, 12. doi: 10.3389/fimmu.2021.730825.

34. Jang, M.J., Coughlin, G.M., Jackson, C.R., Chen, X., Chuapoco, M.R., Vendemiatti, J.L., Wang, A.Z., and Gradinaru, V. (2023). Spatial transcriptomics for profiling the tropism of viral vectors in tissues. Nature Biotechnology, 41(9):1272–1286. doi: 10.1038/s41587-022-01648-w.

35. Ohara, S. (2009). Dual transneuronal tracing in the rat entorhinal-hippocampal circuit by intracerebral injection of recombinant rabies virus vectors. Frontiers in Neuroanatomy, 3. doi: 10.3389/neuro.05.001.2009.

36. Levy, R.B. and Reyes, A.D. (2012). Spatial profile of excitatory and inhibitory synaptic connectivity in mouse primary auditory cortex. Journal of Neuroscience, 32(16):5609–5619. doi: 10.1523/JNEUROSCI.5158-11.2012.

37. Campagnola, L., Seeman, S.C., Chartrand, T., Kim, L., Hoggarth, A., Gamlin, C., Ito, S., Trinh, J., Davoudian, P., Radaelli, C., et al. (2022). Local connectivity and synaptic dynamics in mouse and human neocortex. Science, 375(6585):eabj5861. doi: 10.1126/science.abj5861.

38. Douglas, R.J. and Martin, K.A.C. (2004). Neuronal circuits of the neocortex. Annual Review of Neuroscience, 27. doi: 10.1146/annurev.neuro.27.070203.144152.

39. Patiño, M., Rossa, M.A., Lagos, W.N., Patne, N.S., and Callaway, E.M. (2024). Transcriptomic cell-type specificity of local cortical circuits. Neuron, 112(23):3851–3866.e4. doi: 10.1016/j.neuron.2024.09.003.

40. Pfeffer, C.K., Xue, M., He, M., Huang, Z.J., and Scanziani, M. (2013). Inhibition of inhibition in visual cortex: the logic of connections between molecularly distinct interneurons. Nature Neuroscience, 16(8):1068–1076. doi: 10.1038/nn.3446.

41. Wang, Q. and Burkhalter, A. (2007). Area map of mouse visual cortex. The Journal of Comparative Neurology, 502(3):339–357. doi: 10.1002/cne.21286.

42. Kalatsky, V.A. and Stryker, M.P. (2003). New paradigm for optical imaging: temporally encoded maps of intrinsic signal. Neuron, 38(4):529–545. doi: 10.1016/S0896-6273(03)00286-1.

43. Garrett, M.E., Nauhaus, I., Marshel, J.H., and Callaway, E.M. (2014). Topography and areal organization of mouse visual cortex. The Journal of Neuroscience, 34(37):12587–12600. doi: 10.1523/JNEUROSCI.1124-14.2014.

44. Waters, J., Lee, E., Gaudreault, N., Griffin, F., Lecoq, J., Slaughterbeck, C., Sullivan, D., Farrell, C., Perkins, J., Reid, D., et al. (2019). Biological variation in the sizes, shapes and locations of visual cortical areas in the mouse. PLOS ONE, 14(5):1–13. doi: 10.1371/journal.pone.0213924.

45. Kebschull, J.M. and Zador, A.M. (2015). Sources of PCR-induced distortions in high-throughput sequencing data sets. Nucleic Acids Research, 43(21):e143. doi: 10.1093/nar/gkv717.

46. Ballesteros-Yáñez, I., Benavides-Piccione, R., Elston, G.N., Yuste, R., and DeFelipe, J. (2006). Density and morphology of dendritic spines in mouse neocortex. Neuroscience, 138(2):403–409. doi: 10.1016/j.neuroscience.2005.11.038.

47. Markram, H., Lübke, J., Frotscher, M., Roth, A., and Sakmann, B. (1997). Physiology and anatomy of synaptic connections between thick tufted pyramidal neurones in the developing rat neocortex. The Journal of Physiology, 500 (Pt 2)(Pt 2):409–440. doi: 10.1113/jphysiol.1997.sp022031.

48. Osakada, F. and Callaway, E.M. (2013). Design and generation of recombinant rabies virus vectors. Nature Protocols, 8(8):1583–1601. doi: 10.1038/nprot.2013.094.

49. Wang, Y., Bergelson, S., and Feschenko, M. (2018). Determination of lentiviral infectious titer by a novel droplet digital PCR method. Human Gene Therapy Methods, 29(2):96–103. doi: 10.1089/hgtb.2017.198.

50. Kebschull, J.M., Garcia da Silva, P., Reid, A.P., Peikon, I.D., Albeanu, D.F., and Zador, A.M. (2016). High-throughput mapping of single-neuron projections by sequencing of barcoded RNA. Neuron, 91(5):975–987. doi: 10.1016/j.neuron.2016.07.036.

51. Langmead, B. and Salzberg, S.L. (2012). Fast gapped-read alignment with Bowtie 2. Nature Methods, 9(4):357–359. doi: 10.1038/nmeth.1923.

52. Mayerich, D., Abbott, L., and McCormick, B. (2008). Knife-edge scanning microscopy for imaging and reconstruction of three-dimensional anatomical structures of the mouse brain. Journal of Microscopy, 231(1):134–143. doi: 10.1111/j.1365-2818.2008.02024.x.

53. Ragan, T., Kadiri, L.R., Venkataraju, K.U., Bahlmann, K., Sutin, J., Taranda, J., Arganda-Carreras, I., Kim, Y., Seung, H.S., and Osten, P. (2012). Serial two-photon tomography: an automated method for ex-vivo mouse brain imaging. Nature methods, 9(3):255–258. doi: 10.1038/nmeth.1854.

54. Campbell, R. (2020). SainsburyWellcomeCentre/BakingTray: Jan 2020. Zenodo. doi: 10.5281/zenodo.3631610.

55. Campbell, R., Blot, A., and lguerard. (2020). SainsburyWellcomeCentre/StitchIt: Last release of stitching model 1. Zenodo. doi: 10.5281/zenodo.3941901.

56. Tyson, A.L., Rousseau, C.V., Niedworok, C.J., Keshavarzi, S., Tsitoura, C., Cossell, L., Strom, M., and Margrie, T.W. (2021). A deep learning algorithm for 3D cell detection in whole mouse brain image datasets. PLOS Computational Biology, 17(5):e1009074. doi: 10.1371/journal.pcbi.1009074.

57. Qian, X., Harris, K.D., Hauling, T., Nicoloutsopoulos, D., Muñoz-Manchado, A.B., Skene, N., Hjerling-Leffler, J., and Nilsson, M. (2020). Probabilistic cell typing enables fine mapping of closely related cell types in situ. Nature Methods, 17(1):101–106. doi: 10.1038/s41592-019-0631-4.

58. Patiño, M., Lagos, W.N., Patne, N.S., Tasic, B., Zeng, H., and Callaway, E.M. (2022). Single-cell transcriptomic classification of rabies-infected cortical neurons. Proceedings of the National Academy of Sciences, 119(22):e2203677119. doi: 10.1073/pnas.2203677119.

59. Edelstein, A., Amodaj, N., Hoover, K., Vale, R., and Stuurman, N. (2010). Computer control of microscopes using µManager. Current Protocols in Molecular Biology, 92(1):14.20.1– 14.20.17. doi: 10.1002/0471142727.mb1420s92.

60. Pinkard, H., Stuurman, N., Ivanov, I.E., Anthony, N.M., Ouyang, W., Li, B., Yang, B., Tsuchida, M.A., Chhun, B., Zhang, G., et al. (2021). Pycro-Manager: open-source software for customized and reproducible microscope control. Nature Methods, 18(3):226–228. doi: 10.1038/s41592-021-01087-6.

61. Pati, Y., Rezaiifar, R., and Krishnaprasad, P. Orthogonal matching pursuit: recursive function approximation with applications to wavelet decomposition. In Proceedings of 27th Asilomar Conference on Signals, Systems and Computers, pages 40–44 vol.1, November 1993. doi: 10.1109/ACSSC.1993.342465. ISSN: 1058-6393.

62. Padfield, D. (2012). Masked object registration in the Fourier domain. IEEE Transactions on Image Processing, 21(5):2706–2718. doi: 10.1109/TIP.2011.2181402.

63. Carey, H., Pegios, M., Martin, L., Saleeba, C., Turner, A.J., Everett, N.A., Bjerke, I.E., Puchades, M.A., Bjaalie, J.G., and McMullan, S. (2023). DeepSlice: rapid fully automatic registration of mouse brain imaging to a volumetric atlas. Nature Communications, 14(1): 5884. doi: 10.1038/s41467-023-41645-4.

64. Klein, S., Staring, M., Murphy, K., Viergever, M., and Pluim, J. (2010). elastix: a toolbox for intensity-based medical image registration. IEEE Transactions on Medical Imaging, 29(1): 196–205. doi: 10.1109/TMI.2009.2035616.

65. Bogovic, J.A., Hanslovsky, P., Wong, A., and Saalfeld, S. Robust registration of calcium images by learned contrast synthesis. In 2016 IEEE 13th International Symposium on Biomedical Imaging (ISBI), pages 1123–1126, April 2016. doi: 10.1109/ISBI.2016.7493463. ISSN: 1945-8452.

66. Claudi, F., Petrucco, L., Tyson, A.L., Branco, T., Margrie, T.W., and Portugues, R. (2020). BrainGlobe Atlas API: a common interface for neuroanatomical atlases. Journal of Open Source Software, 5(54):2668. doi: 10.21105/joss.02668.

67. Stringer, C. and Pachitariu, M. (2025). Cellpose3: one-click image restoration for improved cellular segmentation. Nature Methods, 22(3):592–599. doi: 10.1038/s41592-025-02595-5.

68. Wolf, F.A., Angerer, P., and Theis, F.J. (2018). SCANPY: large-scale single-cell gene expression data analysis. Genome Biology, 19(1):15. doi: 10.1186/s13059-017-1382-0.

69. Traag, V.A., Waltman, L., and van Eck, N.J. (2019). From Louvain to Leiden: guaranteeing well-connected communities. Scientific Reports, 9(1):5233. doi: 10.1038/s41598-019-41695-z.

70. McInnes, L., Healy, J., and Melville, J. UMAP: Uniform Manifold Approximation and Projection for dimension reduction, September 2020. arXiv:1802.03426 [stat].

71. Herculano-Houzel, S., Watson, C.R., and Paxinos, G. (2013). Distribution of neurons in functional areas of the mouse cerebral cortex reveals quantitatively different cortical zones. Frontiers in Neuroanatomy, 7. doi: 10.3389/fnana.2013.00035.

72. Murakami, T.C., Mano, T., Saikawa, S., Horiguchi, S.A., Shigeta, D., Baba, K., Sekiya, H., Shimizu, Y., Tanaka, K.F., Kiyonari, H., et al. (2018). A three-dimensional single-cell-resolution whole-brain atlas using CUBIC-X expansion microscopy and tissue clearing. Nature Neuroscience, 21(4):625–637. doi: 10.1038/s41593-018-0109-1.

73. Huang, L., Kebschull, J.M., Fürth, D., Musall, S., Kaufman, M.T., Churchland, A.K., and Zador, A.M. (2020). BRICseq bridges brain-wide interregional connectivity to neural activity and gene expression in single animals. Cell, 182(1):177–188.e27. doi: 10.1016/j.cell.2020.05.029.

